# *De Novo* Peptide Sequencing Reveals a Vast Cyclopeptidome in Human Gut and Other Environments

**DOI:** 10.1101/521872

**Authors:** Bahar Behsaz, Hosein Mohimani, Alexey Gurevich, Andrey Prjibelski, Mark F. Fisher, Larry Smarr, Pieter C. Dorrestein, Joshua S. Mylne, Pavel A. Pevzner

## Abstract

Cyclic and branch cyclic peptides (cyclopeptides) represent an important class of bioactive natural products that include many antibiotics and anti-tumor compounds. However, little is known about cyclopeptides in the human gut, despite the fact that humans are constantly exposed to them. To address this bottleneck, we developed the CycloNovo algorithm for *de novo* cyclopeptide sequencing that employs de Bruijn graphs, the workhorse of DNA sequencing algorithms. CycloNovo reconstructed many new cyclopeptides that we validated with transcriptome, metagenome, and genome mining analyses. Our benchmarking revealed a vast hidden cyclopeptidome in the human gut and other environments and suggested that CycloNovo offers a much-needed step-change for cyclopeptide discovery. Furthermore, CycloNovo revealed a wealth of anti-microbial cyclopeptides from food that survive the complete human gastrointestinal tract, raising the question of how these cyclopeptides might affect the human microbiome.

**SIGNIFICANCE:** The golden age of antibiotics was followed by a decline in the pace of antibiotics discovery in the 1990s. The key prerequisite for the resurgence of antibiotics research is the development of a computational discovery pipeline for antibiotics sequencing. We describe such pipeline for cyclic and branch cyclic peptides (cyclopeptides) that represent an important class of bioactive natural products such as antibiotics and anti-tumor compounds. Our CycloNovo algorithm for cyclopeptide sequencing reconstructed many new cyclopeptides that we validated with transcriptome, metagenome, and genome mining analyses. CycloNovo revealed a wealth of anti-microbial cyclopeptides from food that survive the complete human gastrointestinal tract, raising the question of how these cyclopeptides might affect the human microbiome.

## INTRODUCTION

The golden age of antibiotics was followed by a decline in the pace of antibiotics discovery in the 1990s. However, antibiotics and other natural products are again at the center of attention as exemplified by the recent discovery of teixobactin^1^. The key prerequisite for the resurgence of antibiotics research is the development of computational discovery pipelines^2^ such as the Global Natural Products Social (GNPS) molecular networking^3^, Dereplicator^4^, and VarQuest^5^. The GNPS project already accumulated over a billion mass spectra, an untapped resource for discovery of new antibiotics. However, new algorithms are needed for the GNPS project to realize its promise for antibiotics discovery. Currently, the GNPS network is mainly used for identification of previously discovered natural products and their analogs, emphasizing the need for algorithms for discovery of novel natural products.

This study focuses on *cyclopeptides*, an important class of bioactive natural products with an unparalleled track record in pharmacology: many antibiotics as well as anti-tumor agents, immunosuppressors, and toxins are cyclopeptides. Cyclopeptide sequencing from tandem mass spectra is challenging as their propensity to break at all pairs of points in their cyclic backbone gives a far more complex series of ions than in linear peptides. Cyclopeptides are divided into cyclic *<u>N</u>on-<u>R</u>ibosomal <u>P</u>eptide<u>s</u>* (*NRPs*) and cyclic *<u>Ri</u>bosomally synthesized and <u>P</u>osttranslationally modified <u>P</u>eptide<u>s</u>* (*RiPPs*). NRPs are built from 300 different naturally occurring amino acids according to the complex *non-ribosomal code*^6^ rather than the genetic code. RiPPs are encoded using the genetic code and so built from the twenty proteinogenic amino acids, which however are subjected to numerous post-translational modifications.

The discovery of the cyclopeptide gramicidin S in 1942 (first antibiotic used for treating soldiers during the World War II) led to two Nobel prizes and has been followed by the discovery of ≈400 families of cyclopeptides (*cyclofamilies*) in the last 75 years^5^. A relatively small number of known cyclofamilies reflects the experimental and computational challenges in cyclopeptide discovery. Moreover, the question of how many cyclofamilies remained below the radar of previous studies (even though their spectra have already been deposited to public databases!) remains open.

To answer this question, we consider the problem of recognizing *cyclospectra* (tandem mass spectra that originated from cyclopeptides) that can be matched against biosynthetic genes using various genome mining and peptidogenomics tools^7,8^. These tools typically generate a huge database of putative cyclopeptides, making it prohibitively time-consuming to search large spectral datasets against such databases. Fast algorithms for recognizing cyclospectra are critical for genome mining as they greatly reduce the set of spectra that need to be matched against databases of putative cyclopeptides.

Bandeira et al.^9^ introduced the concept of *spectral networks* that reveal the spectra of related peptides without knowing their amino acid sequences. Nodes in a spectral network correspond to spectra while edges connect *spectral pairs*, i.e. spectra of peptides differing by a single modification or a mutation (see Supplementary Note “Surugamide spectral network.”) Ideally, each connected component of a spectral network corresponds to a cyclofamily^10^ representing a set of similar cyclopeptides. Although spectral networks of various GNPS datasets have become the workhorse of the cyclopeptide studies, they typically contain false positive edges that make analysis of cyclofamilies challenging^3^. Moreover, constructing the spectral network of all GNPS spectra remains an open algorithmic problem.

Recognition of cyclospectra creates a possibility to construct a small *cyclospectral sub-network* of the entire GNPS network and to evaluate the number of cyclofamilies in the GNPS network as the number of connected components in this sub-network. This analysis revealed that many cyclopeptides evaded detection in previous studies and that the known cyclopeptides represent the tip of the iceberg of cyclopeptides that are waiting to be decoded from the GNPS network.

We distinguish between *cyclopeptide identification* (identifying cyclopeptides by matching their spectra against databases of known cyclopeptides^11^) and *de novo cyclopeptide sequencing* (determining the cyclopeptide sequence from a spectrum alone). Although recent studies have made progress towards cyclopeptide identification^4,5,12,13^, the previously developed cyclopeptide sequencing algorithms^14–17^ are rarely used^11^ because they are rather inaccurate, too slow for analyzing high-throughput spectral datasets, and cannot distinguish cyclopeptides from other compounds.

Here we describe CycloNovo, a fast cyclopeptide sequencing algorithm based on the concept of the *de Bruijn graph* of a spectrum, a compact representation of putative *k*-mers (strings formed by *k* consecutive amino acids) in an unknown cyclopeptide (see Supplementary Note “An example of the de Bruijn graph.”) Although de Bruijn graphs represent the workhorse of DNA sequencing^18^, they have not previously been applied to cyclopeptide sequencing. We demonstrate that CycloNovo enables high-throughput analysis of cyclopeptides in large spectral datasets and sequence many cyclopeptides in diverse samples that include marine, soil, and human gut bacterial communities.

### RESULTS

To illustrate how CycloNovo works, we used a spectrum of the cyclopeptide surugamide A^19^(referred to as surugamide hereon) with the amino acid sequence AIIKIFLI (Figure 1).

**Figure 1.**
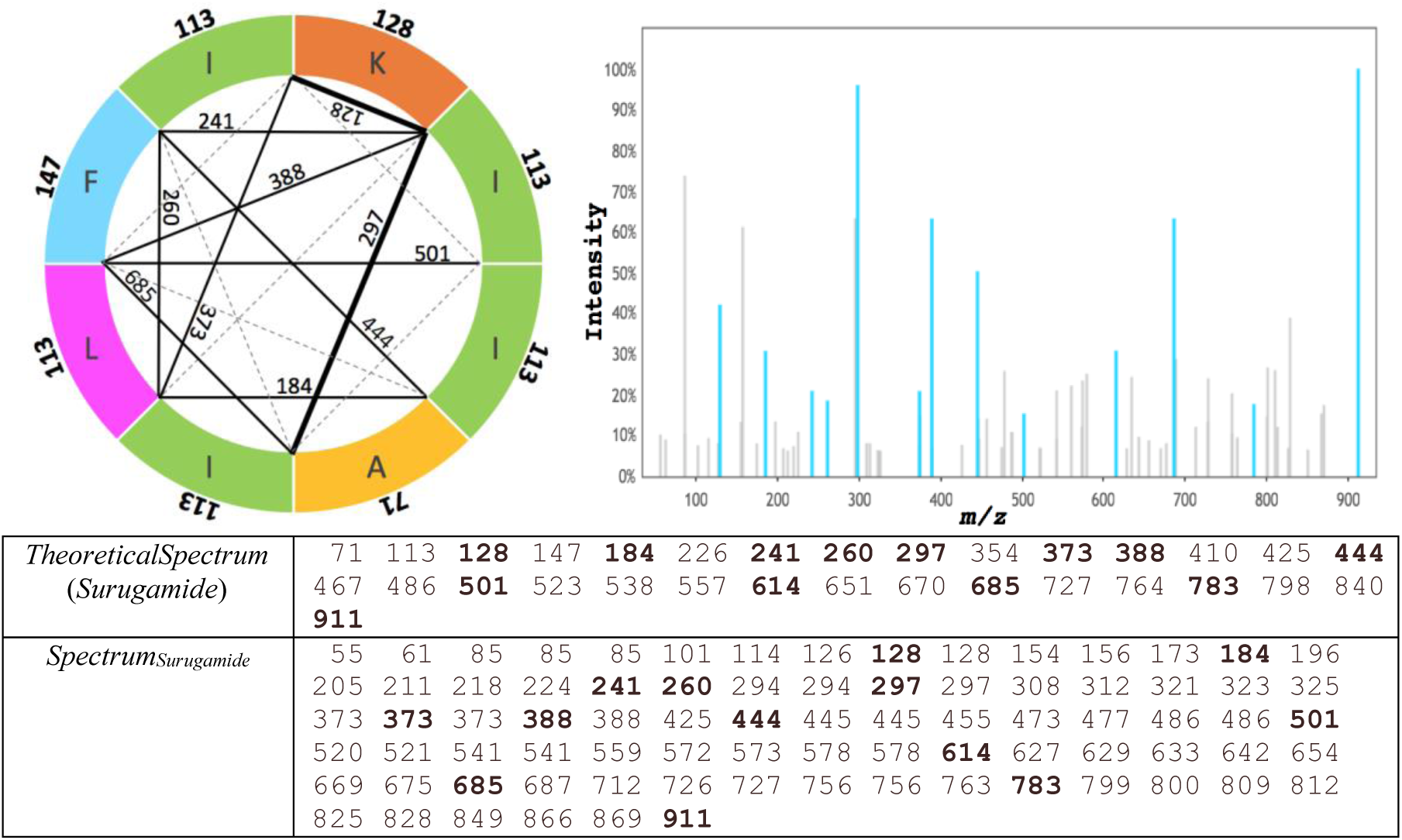
Theoretical and experimental spectra of surugamide. (Top left) Diagram of the surugamide from a marine *Streptomyces* CNQ329 (mass 911.62 Da). Each color represents an amino acid (the numbers on the outer edge are the nominal masses of amino acids in Daltons).

Each chord corresponds to a fragment of surugamide. The solid chords represent the fragments in *TheoreticalSpectrum*(*Surugamide*) whose masses match masses in the experimental spectrum *Spectrum*_*Surugamide*_. The numbers on solid chords show the nominal masses of the corresponding fragment, e.g., the chord labeled 297 corresponds to the fragment Ile-Ile-Ala of mass 297 Da. Each chord corresponds either to a single mass *x* or two masses *x* and *mass(Spectrum)-x* (in the latter case the chord is shown in bold). Given a set of fragments with the same mass, we show one of them (arbitrarily chosen) by a solid chord and the others by dashed chords. For example, one of two fragments with the same integer mass 241 (Ile-Lys and Lys-Ile in clockwise order), is shown by a solid chord and another by a dashed chord. (Top right) The experimental spectrum of surugamide (*Spectrum*_*Surugamide*_) with 82 peaks (GNPS ID MSV000078839). The *y*-axis in the *Spectrum*_*Surugamide*_ shows the ion intensities as the percentage of the intensity of the highest intensity peak. Blue peaks represent masses shared with *TheoreticalSpectrum*(*Surugamide*) for the error threshold *ε* = 0.015 Da. (Bottom) The theoretical and (pre-processed) experimental spectra for surugamide rounded to the nearest integer (this rounding results in the repetitive integers in the list). Masses in the pre-processed experimental spectrum are reduced by the mass of hydrogen *m*_H_≈1.0078 Da. A mass in the theoretical spectrum is shared with a mass in the experimental spectrum if they are within the error threshold. The numbers in bold represent 13 shared masses.

### Theoretical and experimental spectra

Given an amino acid string, its *mass* is defined as the sum of masses of its amino acids. Given a cyclopeptide *Peptide*, its *theoretical spectrum TheoreticalSpectrum*(*Peptide*) is the set of masses of all substrings of *Peptide* (Figure 1). For example, *TheoreticalSpectrum*(AGCD) contains masses of A, G, C, D, AG, GC, CD, DA, AGC, GCD, CDA, DAG, and AGCD. Note that if multiple fragments have the same mass, they contribute a single mass to the theoretical spectrum.

An *experimental spectrum* is a list of *peak*s, where each peak is characterized by its *intensity* and *m/z* (*m* and *z* represent the mass and the charge of the ion corresponding to the peak). For simplicity, we will represent a pre-processed spectrum as an increasing sequence of numbers *Spectrum*={*s*_1_,…, *s*_n_}, assuming that all peaks in the spectrum have charge 1 and ignoring intensities (see Supplementary Note: “Preprocessing spectra.”) We estimate the *PeptideMass* of the cyclopeptide that generated *Spectrum* based on the precursor mass and the charge of *Spectrum.* We define the *symmetric* version of *Spectrum* (denoted *Spectrum\**) as a spectrum that, in addition to all masses in *Spectrum*, contains *PeptideMass-s* for each mass *s* in *Spectrum.*

### Scoring Peptide-Spectrum Matches

A mass *s* in a (pre-processed) experimental spectrum *Spectrum* matches a mass *s’* in *TheoreticalSpectrum*(*Peptide*) if *s* is “equal” to *s’*. By “equal” we mean “approximately equal” with error below the *error threshold ε* (all default values are specified in the Supplementary Note “CycloNovo parameters.”) The score between *Peptide* and *Spectrum* (denoted *score*(*Peptide, Spectrum*) is defined as the number of matches between masses in *Spectrum* and masses in *TheoreticalSpectrum*(*Peptide*). Although CycloNovo uses accurate masses, examples below use nominal masses for simplicity.

Figure 1 illustrates that *score*(*Surugamide, Spectrum*_*Surugamide*_)=13. For a linear peptide *Peptide, score*(*Peptide, Spectrum*) is the number of matches between masses of all linear substrings of *Peptide* and all masses in *Spectrum.* For example, *score*(ILFIK, *Spectrum*_*Surugamide*_)=7 because the theoretical spectrum of the linear peptide ILFIK has 7 shared masses with *Spectrum*_*Surugamide*_corresponding to 7 chords within the ILFIK segment in Figure 1. These chords correspond to the following substrings: K (nominal mass 128), IL (226), IK (241), LF (260), LFI (373), LFIK (501), and ILFIK (614).

### Cyclopeptidic amino acids

Gurevich et al.^5^ combined all currently known peptidic natural products into a single *PNPDatabase*. This database contains 1,257 cyclopeptides (387 cyclofamilies) that we refer to as *CyclopeptideDatabase*. We formed the set of *cyclopeptidic amino acids* (i.e., amino acids that occur in many cyclopeptides) by considering 25 most frequent amino acids in cyclopeptides from *CyclopeptideDatabase* and extending this list to include all proteinogenic amino acids and some common amino acids in RiPPs (see Supplementary Note “Cyclopeptidic amino acids”). 255 out of 1,257 cyclopeptides in the *CyclopeptideDatabase* include only cyclopeptidic amino acids.

### Spectral convolution

The *convolution* of a spectrum is the set of all pairwise differences between its masses^15^. Given a mass *a*, the convolution of *Spectrum with offset a* (denoted *convolution*(*Spectrum, a*)) is defined as the number of masses in the convolution equal to *a* (with error up to *ε*). As shown by Ng et al.^15^, the value *convolution*(*Spectrum, a*) is expected to be high if *a* is the mass of an amino acid in a cyclopeptide that gave rise to *Spectrum*. Thus, offsets with high convolutions reveal the masses of amino acids in an unknown cyclopeptide that gave rise to an experimental spectrum.

To account for measurement errors, we cluster the masses in the convolution using *single linkage clustering* by combining pairs of masses in a cluster if they are less than *ε* apart. We define the *cluster mass* as the median mass of its members, and *cluster multiplicity* as the number of elements in the cluster. We call a cluster *cyclopeptidic* if one of its elements is within of the mass of a cyclopeptidic amino acid. Since high-multiplicity clusters reveal amino acids in the unknown cyclopeptide that gave rise to an experimental spectrum, we use them to generate the set of putative amino acids in an unknown cyclopeptide^15^. See Supplementary Note “Analyzing spectral convolution” for more details.

### CycloNovo outline

Given an experimental spectrum *Spectrum*, the Cyclopeptide Sequencing Problem refers to finding a cyclopeptide *Peptide* that maximizes *score*(*Peptide, Spectrum*). Figure 2 illustrates the CycloNovo pipeline for solving this problem:

- ***Recognizing cyclospectra.*** Natural product researchers use *Marfey’s analysis* for inferring the amino acid composition of an unknown peptide. However, since Marfey’s analysis requires a purified peptide and has a number of limitations^20^, we describe its *in silico* alternative for deriving an *approximate* amino acid composition of a cyclopeptide that gave rise to a given spectrum (see Methods section). If applying this approach reveals that a spectrum originated from a cyclopepeptide, we classify it as a cyclospectrum.
- ***Predicting amino acids in a cyclopeptide.*** For each cyclospectrum, CycloNovo predicts the set of putative amino acids in a cyclopeptide that gave rise to this spectrum. CycloNovo considers each cyclopeptidic cluster with multiplicity exceeding the *cyclopeptidic aa threshold* and classifies the cyclopeptidic amino acids corresponding to this cluster as a *putative amino acid* of the cyclopeptide that generated the cyclospectrum. Figure 2 illustrates that CycloNovo classifies amino acids **A, I/L, F, K,** T, W, R, and G as putative amino acids for *Spectrum*_*Surugamide*_(amino acids occurring in suragamide are shown in bold).
- ***Predicting amino acid composition of a cyclopeptide.*** For each cyclospectrum, CycloNovo uses dynamic programming to find all combinations of putative amino acids with total mass matching the precursor mass of the spectrum. We refer to each such combination as *putative composition Composition(Spectrum)*, which may include the same amino acid multiple times. Figure 2 illustrates that CycloNovo predicts the following putative compositions for *Spectrum*_*Surugamide*_: **A^1^I/L^5^K^1^F^1^ (71^1^113^5^128^1^147^1^)**, I/L^4^F^1^R^2^(113^4^147^1^156^2^), A^2^T^1^K^4^R^1^ (71^2^101^1^128^4^156^1^), and G^1^T^1^I/L^1^K^5^(57^1^101^1^113^1^128^5^). The putative composition of surugamide with one A, five I/L, one K, and one F is represented as **A^1^I/L^5^K^1^F^1^** and is shown in bold.
- ***Predicting k-mers in a cyclopeptide.*** For each *Composition(Spectrum*), it analyzes all linear *k*-mers formed by amino acids in this composition (the default value *k*=5) and scores them against *Spectrum\** using linear scoring. It assumes that if a Peptide-Spectrum Match has a high score *score*(*Peptide,Spectrum*) (a condition that usually holds for well-fragmented spectra), then each linear *k*-mer in *Peptide* also has high score (for an appropriately chosen *k*). High-scoring *k*-mers (defined as *k*-mers with scores exceeding the *k-mer score threshold*) represent putative *k*-mers in an unknown cyclopeptide. For example, for *Composition*=71^1^113^5^128^1^147^1^, there exist 4^5^=1024 5-mers and CycloNovo identifies 524 of them as high-scoring 5-mers. We refer to the set of high-scoring *k*-mers as *Kmers*_*Composition,k*_(*Spectrum*). Figure 2 illustrates that three out of six highest scoring 5-mers for *Spectrum*_*Surugamide*_are correct, i.e., represent 5-mers from surugamide. CycloNovo computes the *k-merScore*, the score of the highest-scoring *k*-mer.
- ***Constructing the de Bruijn graph of a spectrum.*** Given a set *Kmers= Kmers*_*Composition,k*_(*Spectrum*), CycloNovo constructs the *de Bruijn graph* DB_*Kmers*_(*Spectrum*)^18^. Nodes in DB_*Kmers*_(*Spectrum*) correspond to all (*k-1*)-mers from *Kmers* and each directed edge corresponds to a *k*-mer from *Kmers* and connects its first (*k-1*)-mer with its last (*k-1*)-mer. Each cycle in DB_*Kmers*_(*Spectrum*) spells out a cyclic amino acid sequence. Figure 2 presents the *pruned de Bruijn graph* for the putative composition 71^1^113^5^128^1^147^1^ that is obtained by iterative removal of tips (nodes without outgoing or incoming edges), and single isolated edges from the de Bruijn graph. The composition 113^5^128^1^71^1^147^1^ results in a de Bruijn graph with 202 vertices and 524 edges and the pruned de Bruijn graph with 126 vertices and 392 edges (Figure 2).
- ***Generating cyclopeptide reconstructions.*** A cycle in the de Bruijn graph of a spectrum is *feasible* if it spells a cyclopeptide with the mass matching the precursor mass of the spectrum. Using the breadth-first search algorithm, CycloNovo finds all feasible cycles in the de Bruijn graph with length equal to the number of amino acids in *Composition* (a cycle may traverse the same edge multiple times). Each such cycle spells a putative cyclopeptide and CycloNovo scores each of them against *Spectrum.* Finally, it reports the highest scoring cyclopeptides along with the P-values of their Peptide-Spectrum Matches (PSMs) computed using MS-DPR^21^.

**Figure 2.**
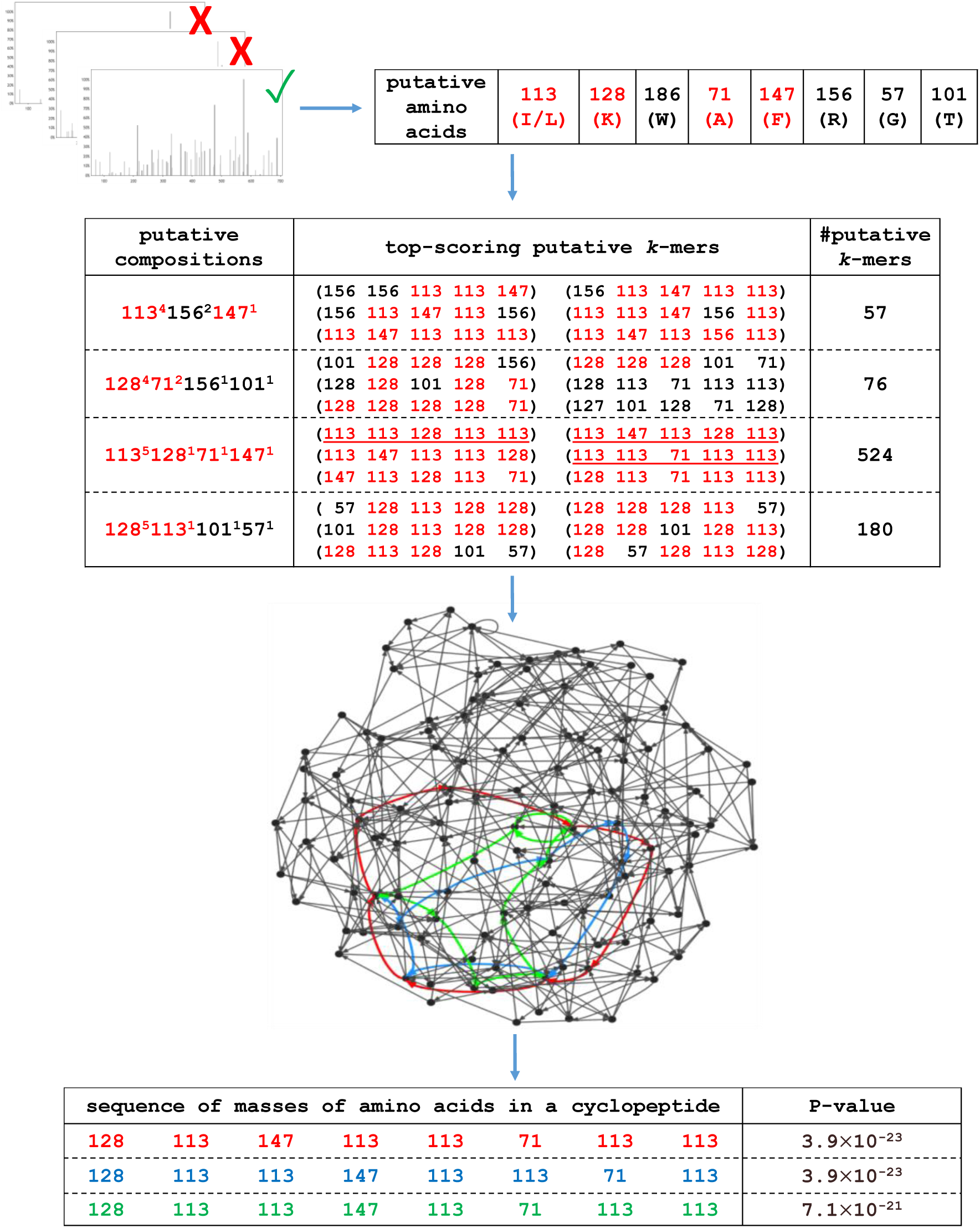
CycloNovo outline illustrated using *Spectrum*_*Surugamide*_. CycloNovo includes six steps: (1) recognizing cyclospectra, (2) predicting amino acids in a cyclopeptide, (3) predicting amino acid composition of a cyclopeptide, (4), predicting *k*-mers in a cyclopeptide, (5) constructing the de Bruijn graph of a spectrum, and (6) generating cyclopeptide reconstructions. Only six top-scoring putative *k*-mers for each putative amino acid composition are shown. Masses of amino acids occurring in surugamide are shown in red and *k*-mers occurring in surugamide are underlined. To simplify the de Bruijn graph (corresponding to the composition 71^1^113^5^128^1^147^1^), all tips and isolated edges in the graph were removed. Red, blue and green feasible cycles in the graph spell out three cyclopeptides shown in the bottom table along with their P-values. The red cycle spells out surugamide.

In the case of *Spectrum*_*Surugamide*_, CycloNovo found three similar cyclopeptides (Figure 2) spelled by feasible cycles in the de Bruijn graph with a putative composition 113^5^128^1^71^1^147^1^ (the highest-scoring one corresponds to surugamide). The remaining three putative compositions do not yield feasible cycles in their de Bruijn graphs (Supplementary Note “De Bruijn Graphs for *Spectrum*_*Surugamide*_.”) CycloNovo sequenced *Spectrum*_*Surugamide*_in ≈3 seconds on a laptop with a single 2.5GHz processor (see Supplementary Note “CycloNovo running time”).

### Datasets

We analyzed various spectral datasets obtained from diverse bacterial communities (Table 1 and Supplementary Note “Information about spectral datasets”). To benchmark CycloNovo, we also analyzed a plant spectral dataset that had a paired RNA-seq dataset, thus enabling us to validate the CycloNovo reconstructions by matching them against the transcriptome.

**Table 1.** Information about various high-resolution spectral datasets analyzed by CycloNovo. The number of distinct cyclopeptides and cyclofamilies was estimated using MS-Cluster^28^ and SpecNets^3^, respectively. The last column shows the number of known cyclopeptides/cyclofamilies (identified by Dereplicator) in each dataset. For each identified cyclopeptide in the CYCLOLIBRARY dataset, we selected the PSM with the minimum P-value (among all PSMs for that cyclopeptide), resulting in a spectral dataset CYCLOLIBRARY with 81 spectra.

| dataset | #spectra | #spectra after preprocessing | #cyclospectra | #distinct cyclopeptides/<br>cyclofamilies | #known cyclopeptides/<br>cyclofamilies |
| --- | --- | --- | --- | --- | --- |
| CYCLOLIBRARY | 81 | 81 | 45 | 45/27 | 45/27 |
| S.VULGARIS | 667 | 212 | 23 | 12/9 | 4/4 |
| HUMANSTOOL | 1,242,178 | 451,962 | 703 | 79/69 | 7/5 |
| GNPS | 51,220,679 | 27,883,895 | 12,004 | 512/213 | 67/37 |
| GNPS<sub>ACTI</sub> | 5,903,921 | 4,435,893 | 1,478 | 116/56 | 38/24 |
| GNPS<sub>CYANO</sub> | 23,582,408 | 12,118,482 | 317 | 74/35 | 5/4 |
| GNPS<sub>PSEUDO</sub> | 697,812 | 581,012 | 2,076 | 120/39 | 5/2 |

The CYCLOLIBRARY dataset contains 81 spectra from 81 distinct cyclopeptides (forming 41 cyclofamilies) that were identified by Dereplicator^4^ after searching the GNPS network against *CyclopeptideDatabase*^5^.

The S.VULGARIS dataset is generated from a single sample collected from seeds of the plant *Senecio vulgaris* (both medicinal and poisonous)^22^ from the *Asteraceae* family. We also analyzed the RNA-Seq reads from the same sample (∼74 million 100 bp long Illumina reads)^22^, assembled them using rnaSPAdes^23^, and used the assembled transcripts (61.9 Mb total length) and prior knowledge of cyclopeptide processing^24–26^ to validate the reconstructed cyclopeptides.

The HUMANSTOOL dataset is generated from 65 stool samples of a single person (L.S., co-author of this paper and a contributor to the “Quantified self” initiative) collected over a course of four years. This dataset is accompanied by the detailed medical and food metadata^27^ as well as metagenomics reads generated from the same samples (project ID <u>PRJEB24161</u>).

The GNPS dataset is formed by combining forty datasets from GNPS^3^. The GNPS_CYANO,_GNPS_PSEUDO,_and GNPS_ACTI_datasets represent sub-datasets of the GNPS dataset corresponding to three phyla with extensively analyzed cyclopeptides (*Cyanobacteria, Pseudomonas* and *Actinomyces*).

### Analyzing the CYCLOLIBRARY dataset

As the cyclopeptides that gave rise to the spectra in the CYCLOLIBRARY dataset are known, we used this dataset to benchmark CycloNovo. We considered a cyclopeptide/spectrum as correctly sequenced if the sequence of the cyclopeptide appeared among reconstructions with three highest-scores. CycloNovo recognized 45 spectra in the CYCLOLIBRARY dataset as cyclospectra and correctly sequenced 38 of these cyclospectra (see Supplementary Note “CycloNovo analysis of the CYCLOLIBRARY dataset”).

### Analyzing the S.VULGARIS dataset

23 recognized cyclospectra in this dataset correspond to twelve distinct cyclopeptides. CycloNovo sequenced ten of them with P-values below 10^−15^(Table 2). Nine of ten reconstructed cyclopeptides match the assembled transcriptome. One reconstructed cyclopeptide (with the highest-scoring reconstruction AFLLADV and score 22), does not match the assembled transcriptome but a suboptimal ALFLGLD reconstruction with score 20 does (see Supplementary Note “Cyclopeptide-encoding transcripts in the S.VULGARIS dataset”).

**Table 2.**
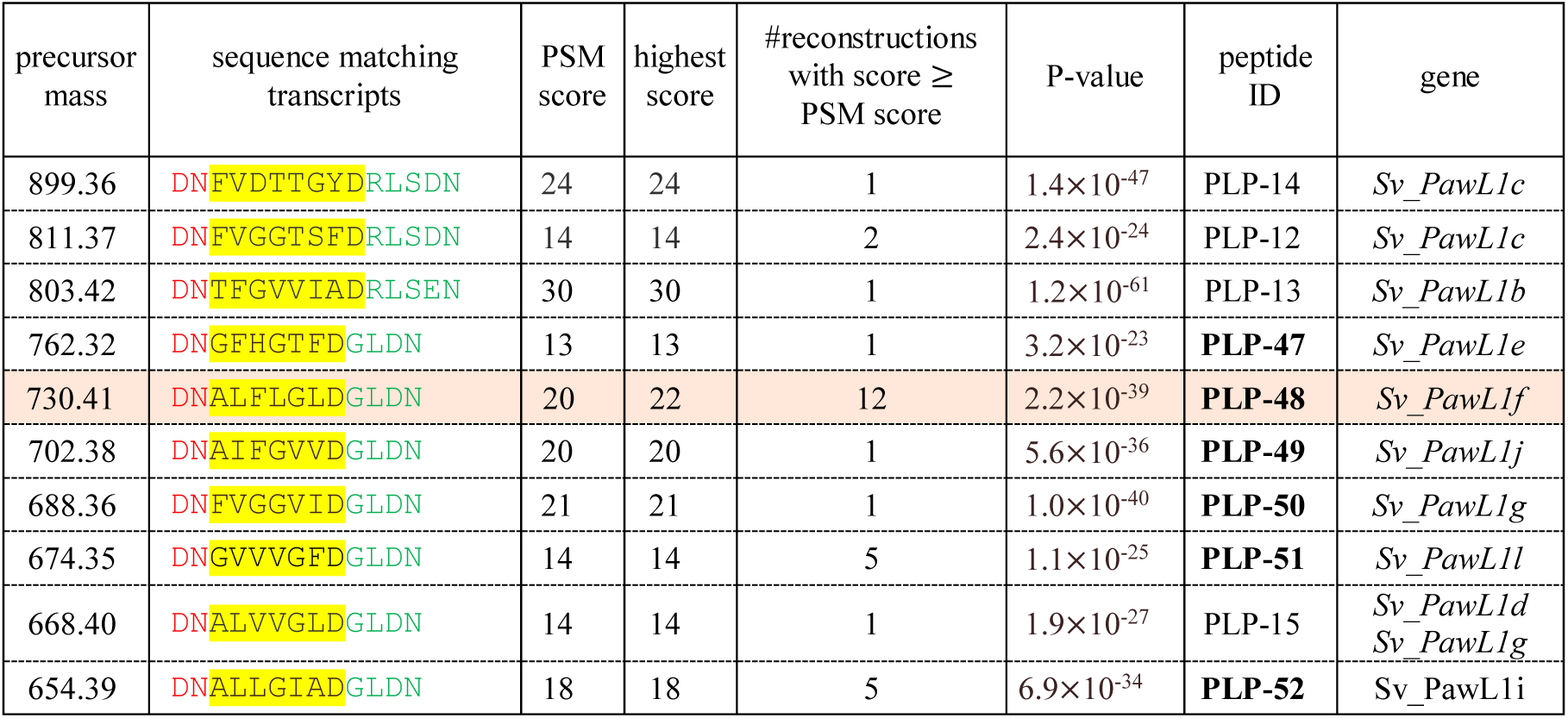
Cyclopeptides reconstructed in the S.VULGARIS dataset. Ten reconstructed cyclopeptides (highlighted in yellow) along with their flanking sequences in transcripts translated into amino acids. For each of these cyclopeptides (reconstructed with P-values below 10-^15^), we selected one representative spectrum with the highest score. The conserved flanking amino acids in the transcripts on the left and right sides of the highlighted cyclopeptides (preceding and succeeding motifs) are shown in red and green, respectively. For nine out of ten cyclopeptides, the reconstruction with the highest score matches one of the transcripts. For the cyclopeptide with mass 730.41 (highlighted in pink), the highest scoring reconstruction AFLLADV (score 22), does not match the assembled transcriptome but a suboptimal ALFLGLD reconstruction (score 20) does. The novel cyclopeptides discovered by CycloNovo are shown with bold IDs and named PLP-47 through PLP-52. For this dataset we used the error threshold ε=0.015 Da as recommended in Fisher *et al* ^24^.

The ten reconstructed cyclopeptides (nine highest-scoring reconstruction and one suboptimal reconstruction) match 11 transcripts (some transcripts encode multiple cyclopeptides and some cyclopeptides are encoded by multiple transcripts) that belong to cyclopeptide-encoding *PawS1-Like* genes in various *Asteraceae* species^22,24^. While three out of eleven identified *PawL1* ORFs and the four cyclopeptides encoded by them (PLP-12 through PLP-15) have been extensively analyzed in recent studies^22,24^, the remaining eight ORFs represent previously unknown cyclopeptide-encoding genes in *S. vulgaris*. See Supplementary Note “Cyclopeptide-encoding transcripts in the S.VULGARIS dataset.”

### Analyzing the HUMANSTOOL dataset

A Dereplicator search of the HUMANSTOOL dataset against *CyclopeptideDatabase* identified seven PSMs at 0% False Discovery Rate (FDR), namely an antimicrobial orbitide citrusin V found in various *Citrus* species^29,30^ and cyclolinopeptides A^31^, B^32^, C^33^, D^33^, H^33^, and E^33^. Cyclolinopeptides are bioactive flaxseed orbitides from *Linum usitatissimum.* The individual who provided the HUMANSTOOL sample frequently ate flaxseeds because they contain α-linolenic acids. CycloNovo sequenced six flaxseed cyclopeptides from the *CyclopeptideDatabase* as well cyclolinopeptide P (a recently discovered cyclopeptide^34^ that has not been added to *CyclopeptideDatabase* yet) as the highest-scoring reconstructions (Supplementary Note “Cyclopeptides in the HUMANSTOOL dataset.”)

In addition to the eight reconstructed orbitides, CycloNovo reconstructed 32 cyclopeptides in the HUMANSTOOL dataset with P-values below 10^−15^ forming 26 cyclofamilies (see Supplementary Notes “Cyclopeptides in the HUMANSTOOL dataset”). Figure 3 shows a connected component in the spectral network formed by four novel cyclopeptides in the HUMANSTOOL dataset and illustrates that CycloNovo reconstructions are consistent with the spectral network.

**Figure 3.**
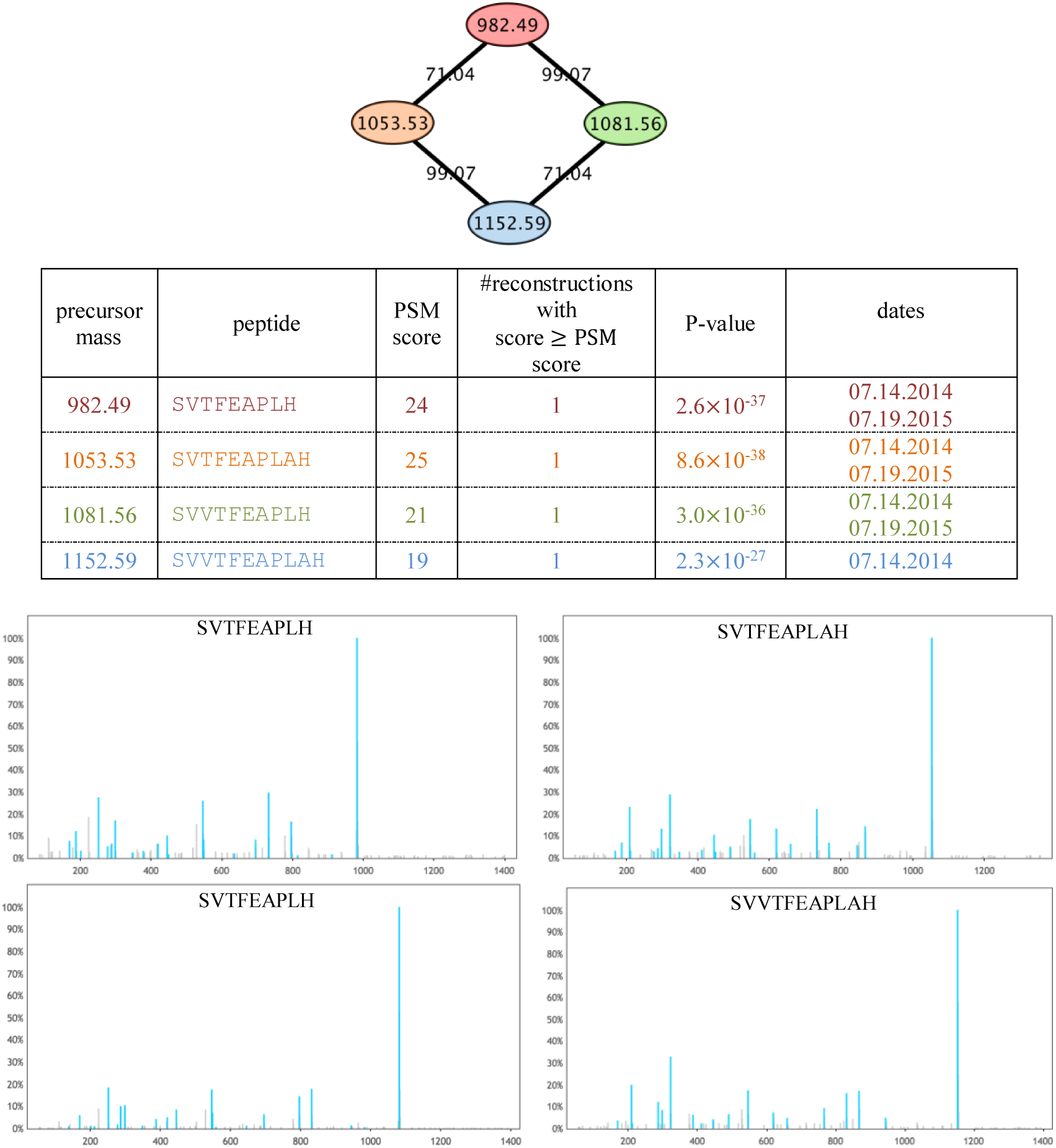
A novel cyclofamily reconstructed by CycloNovo in the HUMANSTOOL dataset. (Top) Four cyclopeptides reconstructed by CycloNovo form a cyclofamily represented by a connected component in the spectral network of the HUMANSTOOL dataset (label “L” stands for one of amino acids L and I). Each node represents a spectrum and two nodes are connected by an edge if their spectral similarity^3^ exceeds 0.8. The numbers on the edges show the mass shifts between the corresponding spectra. (Middle) The *de novo* reconstructions corresponding to the four spectra forming the spectral network. For each cyclopeptide, the cyclic sequence of the highest-scoring reconstruction along with their scores, the number of reconstructions with scores larger or equal to the PSM score (column “#reconstructions with score ≥PSM score”), and P-values are listed. The “dates” column shows the dates when the corresponding samples were taken. Note that the cyclopeptides in this cyclofamily appear on the same dates. (Bottom) The annotated spectra of the four cyclopeptides based on the CycloNovo reconstructions.

The Dereplicator search of all 703 cyclospectra in the HUMANSTOOL dataset against *PNPDatabase* resulted in a single hit and identified a cyclic lipopeptide massetolide F^35^ with P-value 7.5×^−22^10. As this compound includes lipid chains not included in the set of cyclopeptidic amino acids, CycloNovo was not able to generate its full-length reconstructions, but correctly reconstructed its partial amino acid sequence (see Supplementary Note “Cyclopeptides in the HUMANSTOOL dataset.”)

Massetolides are non-ribosomal lipopeptides produced by *Pseudomonas fluoresences*, an indigenous member of human and plant microbiota^36,37^. Analysis of the metagenome assembly of reads paired with the HUMANSTOOL dataset confirmed that *P. fluoresences* is present in the stool samples where massetolide F was detected. Therefore, massetolide F most likely originated from *P. fluoresences* in the human microbiome (see Supplementary Note “Cyclopeptides in the HUMANSTOOL dataset” for information about the assemblies).

### Analyzing the GNPS dataset

We analyzed all cyclospectra in the GNPS dataset using MS-Cluster^28^ and SpecNets^38^ with the goal of estimating the number of still unknown cyclopeptides and cyclofamilies originating from spectra already deposited into GNPS. To provide a conservative estimate for the number of cyclopeptides and cyclofamilies, we limited the analysis to clusters with at least three spectra. 12,004 cyclospectra in the GNPS dataset originated from 512 cyclopeptides and 213 cyclofamilies. Dereplicator search of these cyclospectra against *CyclopeptideDatabase* identified only 67 cyclopeptides from 37 cyclofamilies (see Supplementary Note “Cyclopeptides in the GNPS dataset”). For each putative cyclopeptide, we selected a representative spectrum with the highest *k-merScore*, resulting in 512 spectra corresponding to the 512 cyclopeptides. CycloNovo *de novo* sequenced 94 cyclopeptides with P-values below 10^−15^ in this set of 512 cyclospectra (see Supplementary File).

### Comparing CycloNovo and Dereplicator

Figure 4 compares the number of distinct cyclopeptides, including some branch-cyclic peptides, (see Supplementary Note “Cyclopeptides in the HUMANSTOOL dataset”) and cyclofamilies revealed by CycloNovo and identified by Dereplicator in searches against the *PNPDatabase*. As Figure 4 illustrates, even for the extensively studied phyla of *Cyanobacteria* and *Pseudomonas*, only a small fraction of cyclopeptides and cyclofamilies revealed by CycloNovo are currently known. Moreover, CycloNovo revealed many novel cyclopeptides in known cyclofamilies. For example, CycloNovo reconstructed six novel variants of surugamide by analyzing the GNPS^ACTI^ dataset and revealed the widespread proliferation of the recently described *A-domain skipping* phenomenon^5,39^, suggesting that it is more prevalent than was previously thought (each A-domain encodes a single amino acid in an NRP according to non-ribosomal code). Genome mining efforts typically rule out such events due to the consecutive arrangements of A-domains in NRP synthetases (see Supplementary Note “Surugamide spectral network.”).

**Figure 4.**
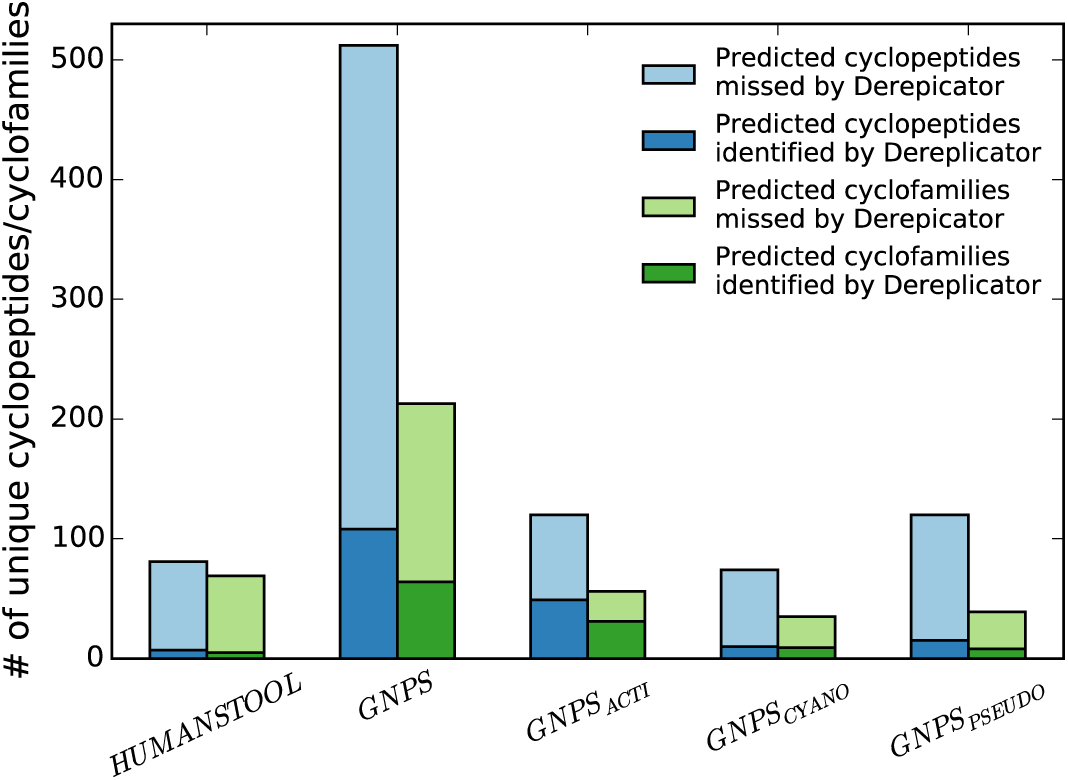
Number of cyclopeptides (blue bars) and cyclofamilies (green bars) predicted by CycloNovo and identified/missed by Dereplicator in various spectral datasets. Missed cyclopeptides/cyclofamilies are not present in *PNPDatabase*.

## DISCUSSION

Although the advent of the GNPS molecular network has created a new resource for natural product discovery, there exists a large body of still unknown bioactive compounds represented by various spectra in the GNPS network^4^ (less than one percent of GNPS spectra have been identified so far). As the existing database search approaches are limited to identifying known cyclopeptides and their variants, *de novo* cyclopeptide sequencing is needed to reveal the “dark matter of cyclopeptidomics.”

Charlop-Powers et al.^40^, recently demonstrated that New York urban parks (rather than exotic areas like rainforests or coral reefs) represent an untapped resource for discovery of clinically important non-ribosomal peptides. This surprising discovery illustrated the still unexplored biosynthetic potential of environmental metagenomes but has not revealed the chemical compounds that these metagenomes encode. As Nothias et al.^41^, wrote in the follow-up commentary *Antibiotic discovery is a walk in the park*, Charlop-Powers et al., revealed the potential for discovering antibiotics in our backyards but did not answer the question what these molecules are and whether they are actually produced by the microbes. To address these questions, we analyzed the GNPS molecular network (our “digital version” of the New York Central Park that contains mass spectra from many environmental samples) and demonstrated that it contains spectra that originated from hundreds of still unknown cyclopeptides.

Only 81 out of 1,257 known cyclopeptides (42 out of 387 known cyclofamilies) have been identified in the GNPS network^5^. CycloNovo revealed over 400 unknown cyclopeptides from 176 novel cyclofamilies by analyzing only ≈51 million GNPS spectra, illustrating that the currently known cyclopeptides represent just a small fraction of cyclopeptides whose spectra have been already deposited into the GNPS network. CycloNovo correctly sequenced many known cyclopeptides in a blind mode and reconstructed many novel cyclopeptides that were validated using transcriptomics data.

Our analysis of the HUMANSTOOL dataset demonstrates that numerous bioactive cyclopeptides from consumed plants remain stable throughout the proteolytic, absorptive and microbial ecosystem provided by the gastrointestinal system and thus interact with human microbiome. It also found cyclospectra originating from the branch cyclic peptide massetolide produced by an indigenous member of the human microbiota and confirmed by metagenomics analysis. In addition, it revealed a large number of still unknown cyclopeptides in the human gut that are either a part of the human diet or are products of the human gut microbiome.

The cyclolinopeptides constitutes the largest identified component of the spectral network of all cyclospectra in the HUMANSTOOL dataset. Controlled diets have demonstrated beneficial effects of flaxseed consumption^32^ and revealed that flaxseed is an effective chemo-preventive agent^42^. However, it remains unknown how flaxseed cyclopeptides affect the human microbiome. Previous studies of flaxseed focused on α-linolenic acid^43^ and other bioactive compounds^44^ affecting the digestive system. However, none of the flaxseed studies have identified what specific flaxseed ingredients are associated with the observed biological outcomes. As discussed in Shim et al.^43^, attributing a certain bioactivity to specific flaxseed compounds is a difficult task as multiple compounds are present in various flaxseed fractions.

It is remarkable that cyclolinopeptides remain stable in a proteolytic environment of the human gut and are not degraded. Although the immunosuppressive potential of cyclolinopeptides was established two decades ago^32^, little was known about their antimicrobial potential. Recent studies demonstrated significant antimicrobial activities of flaxseed, but it remains unclear which specific compounds are responsible for these activities^45^. As our analysis revealed that cyclolinopeptides survive the human digestive tract, we propose that the antimicrobial activities of flaxseeds might be caused by cyclolinopeptides, complementing their known anti-fungal^46^ and anti-malarial^47^ activities. Finding antimicrobial cyclopeptides in human stool raises the question of how these bioactive antimicrobial cyclopeptides might affect the human microbiome.

## METHODS

### Spectral convolution

We represent each spectrum *Spectrum*={*s*_1_,…, *s*_n_} as its *spectral diagram*, the set of *n×*(*n-1*)/*2* 2-dimensional points (*s*_*i*_, *s*_*j*_) for 1 ≤ *i* ≤ *j* ≤ *n*. Given a mass *a*, the convolution of *Spectrum with offset a* (denoted *convolution*(*Spectrum, a*)) is equivalently defined as the number of points in the diagonal (45°) band *y ≈ x+a* in the spectral diagram. Figure 5 presents the spectral diagram of *TheoreticalSpectrum*(AGCD) and reveals that bands corresponding to its amino acids (71, 57, 103, and 115 Da) are the most *populous* (contain a large number of points as compared to other bands), i.e., *convolution*(*Spectrum,a*) is high when *a* is the mass of amino acids A, G, C, or D. For example, *TheoreticalSpectrum*(AGCD) includes five pairs of fragment masses ((G,AG), (D,AD), (AGC,GC), (CD,CDA), and (GCD,AGCD)) that are located on the “blue” diagonal *y = x+mass(A)* in Figure 5.

**Figure 5.**
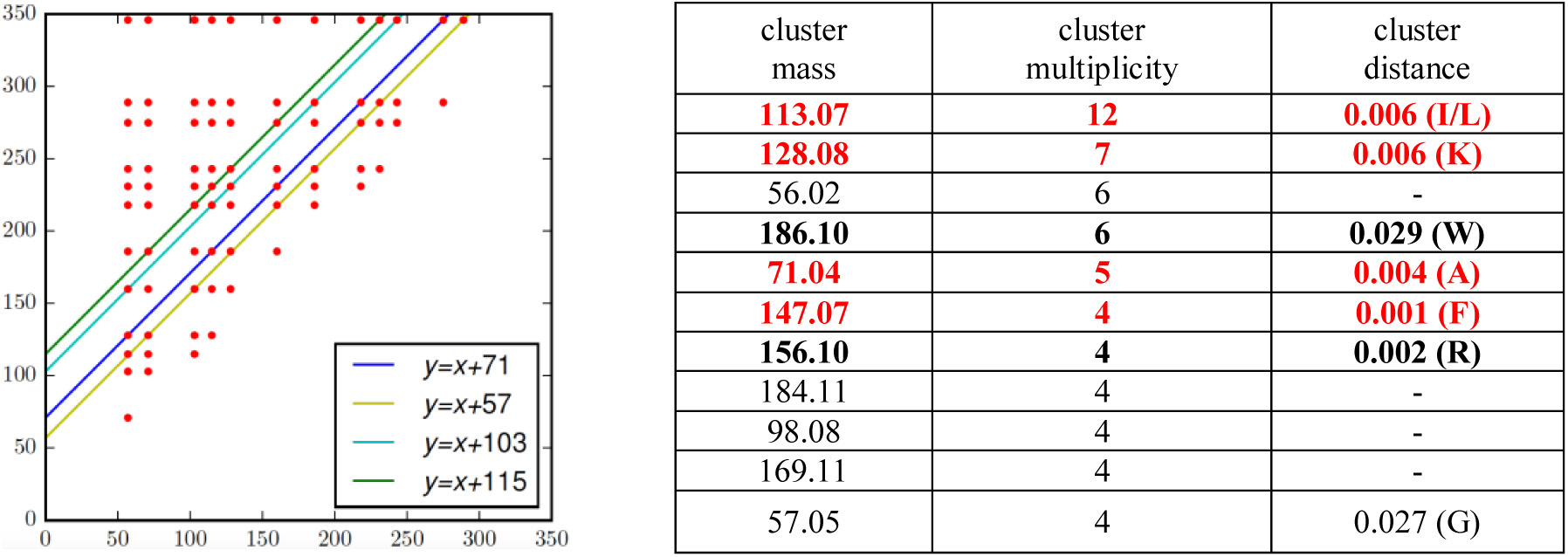
The spectral diagram of *TheoreticalSpectrum*(AGCD) (left) and the list of clusters in the convolutions of *Spectrum*_*Surugamide*_ (right). (Left) The highlighted lines with slope 1 correspond to the masses of the amino acids, A, G, C, and D and contain 5, 9, 5, and 6 points, respectively. (Right) Clusters in the convolutions of *Spectrum*^*Surugamide*^ in the decreasing order of their multiplicities. Only clusters with masses between 55 and 190 Da and multiplicity exceeding 3 are shown. Cyclopeptidic clusters are shown in bold and cyclopeptidic clusters with masses similar to the masses of amino acids in surugamide are shown in red. *Cluster distance* is defined as the distance between the cluster mass and a closest mass of a cyclopeptidic amino acid.

The spectral diagrams for *TheoreticalSpectrum*(*Surugamide*) and experimental *Spectrum*_*Surugamide*_ highlight four populous diagonal bands *y ≈ x+a*, where *a* is the mass of one of four amino acids in surugamide with integer masses 71, 113, 128, and 147 (Supplementary Note “Analyzing spectral convolution.”) These populous bands in the spectral diagram reveal the masses of amino acids in an unknown cyclopeptide that gave rise to an experimental spectrum.

Figure 5 lists high-multiplicity clusters for *Spectrum*_*Surugamide*_(see Supplementary Note “Analyzing spectral convolution”) and shows that many of them have masses that are similar to the masses of amino acids in surugamide. Since populous diagonals (high-multiplicity clusters) in the spectral diagram reveal amino acids in the unknown cyclopeptide that gave rise to an experimental spectrum, we use them to generate the set of putative amino acids^18^.

### Recognizing cyclospectra

A cluster in the spectral convolution is called *frequent* if its multiplicity exceeds the *cluster multiplicity threshold* (the default threshold for *Spectrum*_*Surugamide*_is 7). CycloNovo classifies a spectrum as a cyclospectrum if the number of frequent cyclopeptidic clusters in its spectral convolution is at least *minNumberFrequentClusters* (the default value *minNumberFrequentClusters*=2). Since there exist two frequent cyclopeptidic clusters for *Spectrum*_*Surugamide*_(corresponding to amino acids I/L and K), it is classified as cyclopeptidic (Figure 5). In addition to *Spectrum*_*Surugamide*_, out of 938 spectra passing the preprocessing step in the small spectral dataset for *Streptomyces CNQ329* that contains *Spectrum*_*Surugamide*_, CycloNovo recognized only one cyclospectrum, also originated from surugamide. See Supplementary Note “Distinguishing cyclospectra from spectra of linear peptides and polymers” for selecting CycloNovo parameters.

### Estimating the number of distinct cyclopeptides and cyclofamilies

Spectral datasets often contain multiple spectra originating from the same compound. CycloNovo clusters similar cyclospectra using MS-Cluster^28^ and estimates for the number of distinct cyclopeptides as the number of constructed clusters. It further constructs the spectral network of cyclospectra using SpecNets^3^ and estimates for the number of distinct cyclofamilies as the number of connected components in this network.

## Acknowledgements

We thank Ben Pullman, Sergey Nurk, Alexey Melnik, and Louis-Felix Nothias for fruitful discussions.

## Availability

CycloNovo is available as both a stand-alone tool (https://github.com/bbehsaz/cyclonovo) and a web application (http://gnps.ucsd.edu/ProteoSAFe/static/gnps-theoretical.jsp). All described datasets are available through the corresponding public repositories.

## Funding

The work of B.B., H.M. and P.A.P. was supported by the US National Institutes of Health (grant 2-P41-GM103484). The work of B.B. was also supported by the Natural Science and Engineering Research Council of Canada. The work of A.G. and A.P. was supported by the Russian Science Foundation (grant 14-50-00069). J.S.M. was supported in part by an Australian Research Council Future Fellowship (FT120100013).

M.F.F. was supported by the Australian Government’s Research Training Program and a Bruce and Betty Green Postgraduate Research Scholarship.

Pavel Pevzner is a co-founder, has an equity interest and receives income from Digital Proteomics, LLC. The terms of this arrangement have been reviewed and approved by the University of California, San Diego in accordance with its conflict of interest policies.

### Contributions

P.A.P. and B.B. designed the CycloNovo algorithm and B.B. implemented it. B.B. did the benchmarking and spectral network analysis for all datasets included in this study. M.F. and J.M. generated the S.VULGARIS spectral dataset and helped with the biological interpretation and validation of the identified cyclopeptides. A.P. assembled RNA-seq data for S.VULGARIS. L.S. contributed the HUMANSTOOL sample set for analysis and commented on the results. H.M and A.G. performed the Dereplicator search for all datasets. P.C.D., H.M. and P.A.P. directed the work. B.B. and P.A.P. wrote the manuscript with contributions from all co-authors.

## SUPPLEMENTARY NOTES

### Supplementary Note: Surugamide spectral network

Supplementary Figure S1 shows a connected component in the spectral network containing known and novel surugamide variants (spectral dataset GNPS^ACTI^ generated from samples collected from various *Actinomyces* species).

**Supplementary Figure S1.**
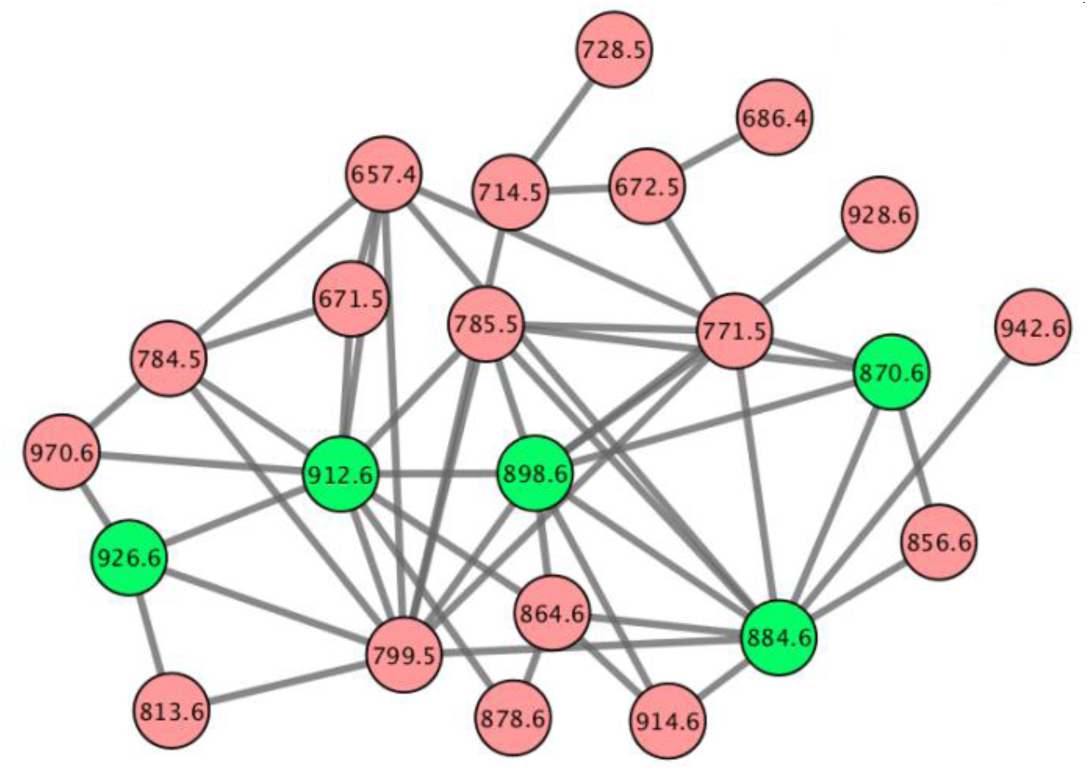
A connected component in the spectral network that contains various surugamide variants. Each node in the network is labeled by the precursor mass of a spectrum and each edge connects spectral pairs that reveal related cyclopeptides. The five green nodes are the known surugamide variants^1^. The pink nodes represent unknown cyclopeptides. The spectral network was constructed based on all cyclospectra in the GNPS_ACTI_ dataset.

Supplementary Figure S2 shows a subgraph of the spectral network shown in Supplementary Figure S1 that includes only known surugamides and six novel variants reconstructed by CycloNovo. Supplementary Figure S3 illustrates that five of these novel variants differ from known surugamides by deletions of some amino acids.

**Supplementary Figure S2.**
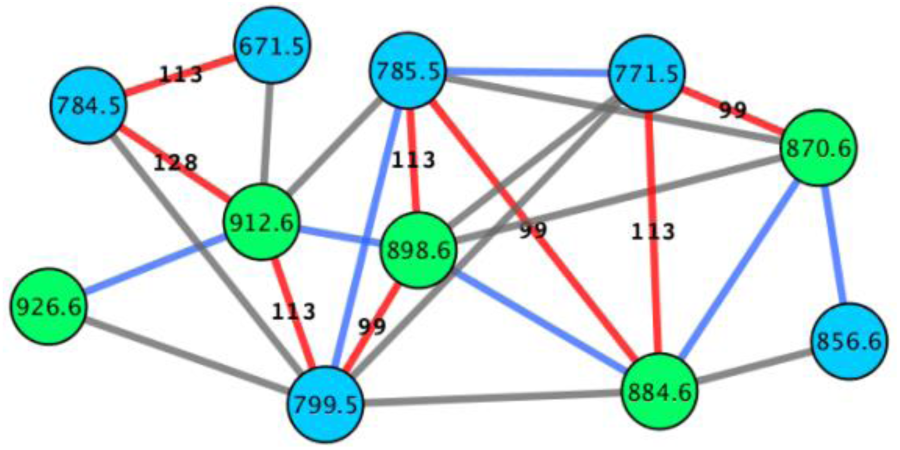
A subgraph of the suragamide connected component in the spectral network of all cyclospectra from the GNPS_ACTI_ dataset showing only the known and novel surugamide variants sequenced by CycloNovo. The green nodes correspond to known surugamides and the blue nodes represent the novel surugamide variants reconstructed by CycloNovo. The numbers on edges represent the nominal mass shift between the corresponding spectra. The red edges highlight the mass shifts that suggest loss/addition of an amino acid in the peptide and the blue edges connects peptides that differ from each other by a single Ile **→** Val or Val **→** Ile substitution (resulting in a nominal offset 14 Da). Although the 14 Da offset can also correspond to methylation, the substitutions represent the more likely explanations in this case. The grey edges show mass shifts that represent combinations of those mass shifts. Supplementary Figure S1 presents the entire connected component.

**Supplementary Figure S3.**
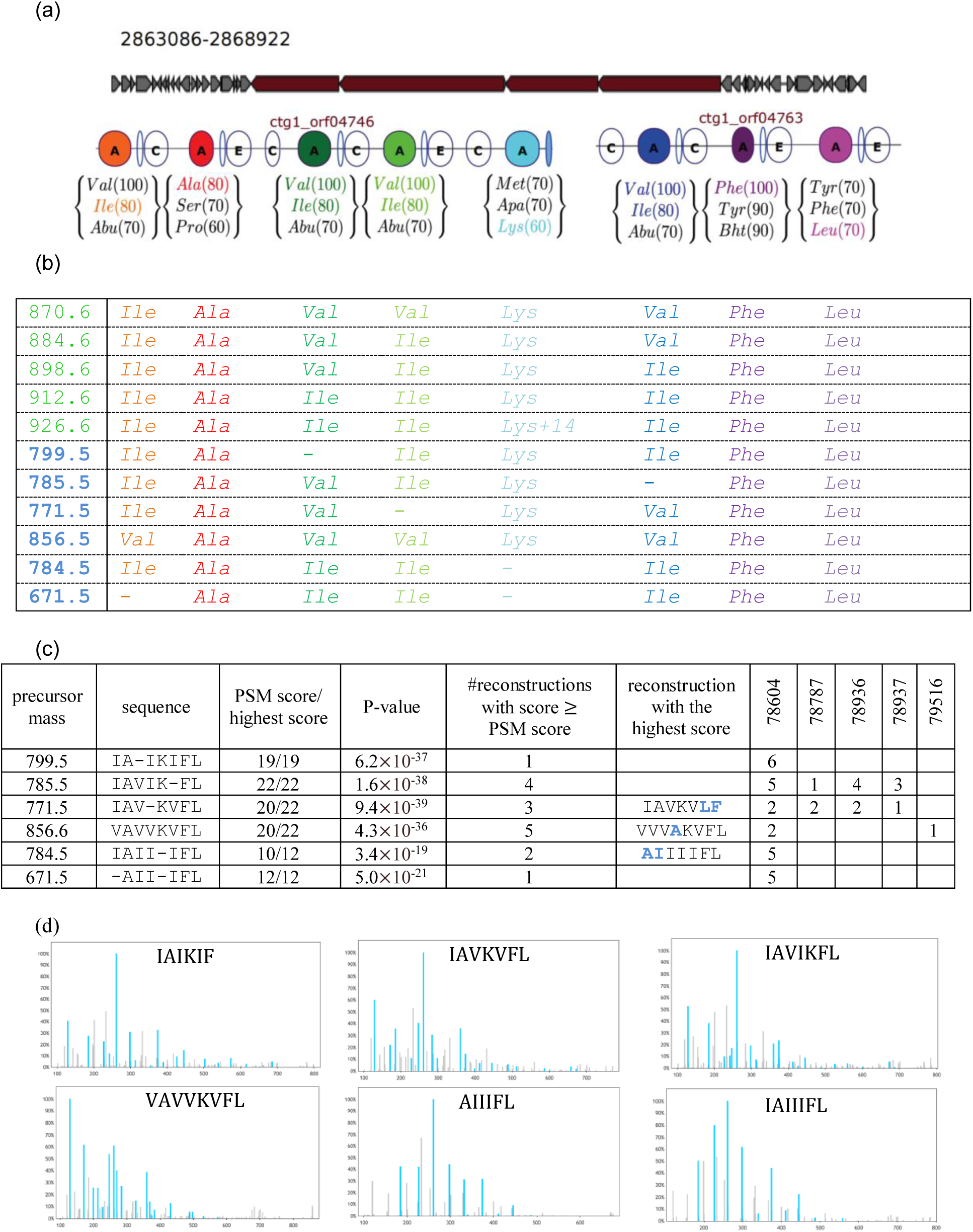
Known and novel surugamide variants. (a) Surugamide gene cluster in *Streptomyces albus* along with the three most likely amino acids for each A-domain and their scores predicted by NRPSpredictor2^2^. See Mohimani et al, 2017^1^ for more details on this representation. (b) Five known (first five rows) and six novel (last six rows) surugamide variants. Each column is color-coded based on the color of the A-domain they represent in the top figure. The dash symbols indicate a violation of the non-ribosomal code (A-domain skipping) when an A-domain in the surugamide gene cluster does not “#reconstructions with score ≥ add an amino acid to a cyclopeptide. (c) *De novo* reconstructions of the novel surugamide variants. The column ‘PSM score/highest score’ shows the score of the cyclopeptide and the highest score observed for that spectrum among all CycloNovo reconstructions. The “P-value” column presents the P-value of the PSM (for each cyclopeptide, the spectrum that yielded the lowest P-value is reflected). The column PSM score” shows the number of reconstructions with score greater or equal to the PSM score. The column “reconstruction with the highest score” shows a highest-scoring reconstruction for the cases when the PSM score is below the highest score. The number of spectra corresponding to each novel surugamide variant in the five GNPS datasets are presented in the columns ‘78604’, ‘78787’, ‘78936’, ‘78937’, and ‘79516’, representing the GNPS sub-datasets MSV000078604, MSV000078787, MSV000078936, MSV000078937, and MSV000079516, respectively. Finding the same surugamide variants in different studies makes it unlikely that they represent artifacts. (d) Annotated spectra of six novel surugamide variants.

### Supplementary Note: An example of the de Bruijn graph

Figure S4 presents an example of the de Bruijn graph.

**Supplementary Figure S4.**
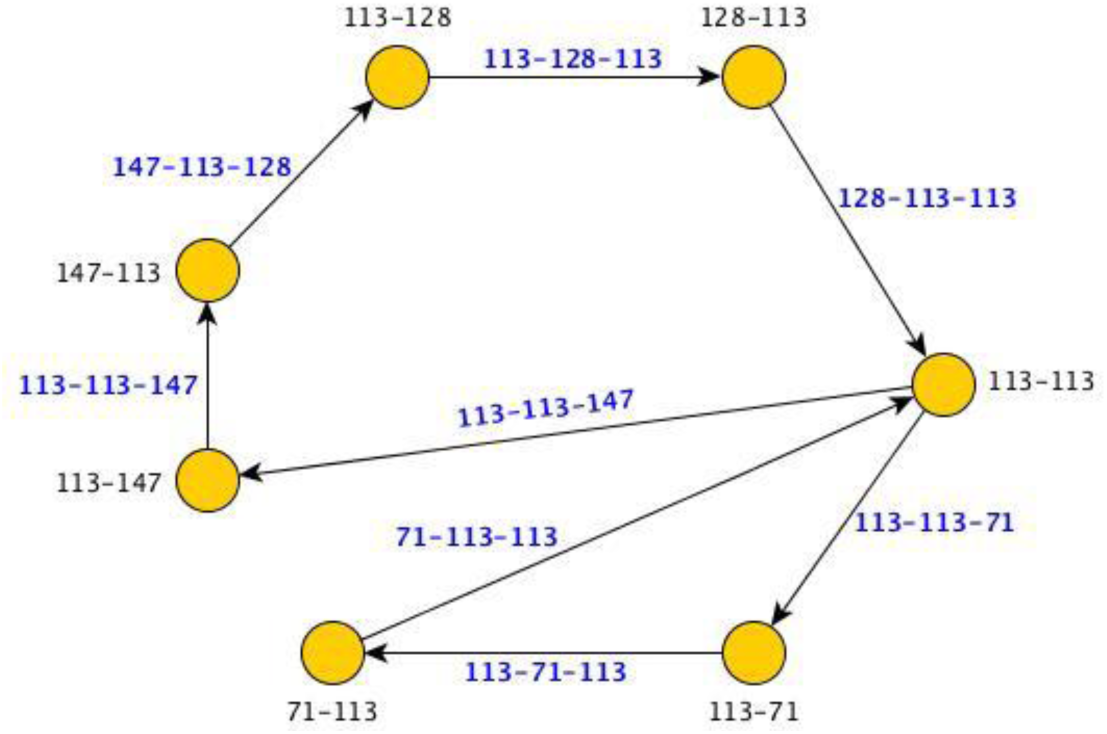
The de Bruijn graph constructed from eight 3-mers of surugamide. The sequence of nominal masses of amino acids in surugamide is represented as 71-113-113-147-113-128-113-113. Nodes (edges) in the de Bruijn graph correspond to seven 2-mers (eight 3-mers) in surugamide. Each edge (3-mer) connects the node corresponding to its initial 2-mer to the node corresponding to its final 2-mer. A traversal of edges of the graph spells out the sequence of masses of amino acids in surugamide.

### Supplementary Note: Preprocessing spectra

Similar to pre-processing practices in proteomics^3^, CycloNovo filters out low-intensity peaks in each spectrum by retaining at most 5 peaks with the highest intensities in each 50 Da window. CycloNovo further filters out all peaks that are less than 0.05 Da apart from another peak with higher intensity. It further removes spectra with a small number of peaks (less than 20) and spectra with a small precursor mass (less than 500 Da). We subtract the mass of a hydrogen atom from all masses in the spectrum (for simplicity, we assume that each ion is protonated with a single proton).

### Supplementary Note: CycloNovo parameters

Table S1 specifies the default values of CycloNovo parameters.

**Supplementary Table S1.** The default values of CycloNovo parameters along with the command-line arguments to specify their values. The parameters introduced in the main text are shown in green rows and the parameters introduced in the Supplementary Note “Distinguishing cyclospectra from spectra of linear peptides and polymers” are shown in pink rows. For all datasets in this paper, all cyclospectra were, recognized with the default parameters α = 0.007 and β = −1 except for cyclospectra in the GNPS datasets that were recognized with even more conservative parameters. α = 0.008 and β = 0

| Universal Parameter | default value | command line argument |
| --- | --- | --- |
| error threshold (Da) | 0.02 | --precursor_ion_thresh |
| Parameters used for recognizing cyclospectra |  |  |
| $\alpha$ | 0.07 | --alpha |
| $\beta$ | -1 | --beta |
| *cycloIntensity* threshold | 60% | --cyclointensity |
| *k-merScore* threshold | 4 | --kmer_score |
| *minNumberFrequentClusters* threshold | 2 | --num_frequent_clusters |
| **Parameters used for *de novo* sequencing** |  |  |
| *cyclopeptidic aa* threshold | 1 | --aminoacid_multiplicity |
| *k-mer* threshold | 2 | --kmer_threshold |
| *k-mer size* | 5 | --kmer_size |

### Supplementary Note: Distinguishing cyclospectra from spectra of linear peptides and polymers

#### The challenge of distinguishing cyclospectra from spectra of linear peptides and polymers

Fragmentation of linear peptides typically results in *prefix* (e.g., b-ions) and *suffix* (e.g., y-ions) ions and rarely generates *internal* ions. However, spectra of some linear peptides feature a substantial number of internal ions, leading to a possibility to erroneously classify them as cyclospectra. Another source of a potential misclassification of some spectra as cyclospectra are polymers that represent a common source of contamination in mass spectral datasets. Since polymers are made up of repeated units, the spectral convolution of a polymer spectrum typically has high-multiplicity clusters (for clusters corresponding to masses of the repeat units). In some cases, the adducts of these repeat units form high multiplicity clusters with masses equal to the masses of a cyclopeptidic amino acid, triggering a possibility to misclassify a polymer spectrum as a cyclospectrum.

#### LINEARLIBRARY and POLYMERLIBRARY datasets

To ensure that CycloNovo does not misclassify spectra of linear peptides and polymers as cyclospectra, we analyzed two spectral datasets described below:

- LINEARLIBRARY is a set of 105,871 Collision-Induced Dissociation (CID) tandem mass spectra of distinct linear peptides from the Massive Knowledge-Based spectral library^4^ of linear peptides distilled from all human proteomics data in the MassIVE database.
- POLYMERLIBARY is a set of 448 tandem spectra generated from polyethylene glycol (MSV000081544).

These spectral datasets have spectra with the precursor masses varying between 500 Da and 2000 Da and the charges at most 2.

#### Additional tests for recognizing cyclospectra

To distinguish cyclospectra from spectra of linear peptides and polymers, CycloNovo only classifies a spectrum as cyclopeptidic if it passes additional tests described below.

- **High multiplicity cyclopeptidic clusters test (distinguishing cyclospectra from spectra of linear peptides)**. As described in the main text, CycloNovo first selects a spectrum for further analysis if its spectral convolution has at least *minNumberFrequentClusters* frequent cyclopeptidic clusters, i.e., clusters with multiplicities exceeding the cluster multiplicity threshold. Since the cluster multiplicities typically increase with the increase in the length of a peptide, this threshold increases with the increase in the peptide mass. We thus defined the *cluster multiplicity threshold* as *α×precursorMass+β* (see below for selecting parameters *α* and *β*).
- **Polymer test (distinguishing cyclospectra from polymer spectra)**. For each cyclospectrum *Spectrum*, CycloNovo analyzes clusters with masses of repeat units observed in background contamination from polyethylene glycol, NaCl, polypropylene glycol, and trimethylsiloxane (44.03, 57.96, 58.04, and 72.04 Da, respectively). We refer to these masses as *polymeric units*^5^ and refer to clusters with masses equal to polymeric units as *polymer-clusters*. CycloNovo classifies a spectrum as polymeric if there exist at least *minNumberFrequentClusters* polymer-clusters with multiplicities at least the *cluster multiplicity threshold.* Polymeric spectra are filtered out from the set of found cyclospectra.
- ***cycloIntensity* test**. For each cyclospectrum, CycloNovo considers all frequent cyclopeptidic clusters. For each such cluster of mass *a*, we consider all pairs of masses *x* and *y* in the spectrum contributing to this cluster, i.e., satisfying the condition *y≈x+a.* The *cyclointensity* of the spectrum, referred to as *cycloIntensity*, is defined as the total intensity of all such peaks (across all frequent cyclopeptidic clusters) divided by the total intensity of all peaks in *Spectrum*. Spectra with cyclointensity below the *cycloiIntensity threshold* are filtered out.
- ***k-merScore* test**. CycloNovo computes the *k-merScore*, the score of the highest-scoring *k*-mer that contributes to the de Bruijn graph of the spectrum and filters out cyclospectra with *k-merScore* below the *k-merScore threshold*.

#### Selecting thresholds for recognizing cyclospectra

To select the default value of *cluster multiplicity threshold*=*α×precursorMass+β*, we varied parameters *α* (from 0.005 to 0.02) and *β* (from −5 to +5) and analyzed all found cyclospectra in the CYCLOLIBRARY, LINEARLIBRARY, and POLYMERLIBARY datasets (Supplementary Figure S5). Despite its smaller size, CYCLOLIBRARY is the only dataset where CycloNovo recognizes cyclospectra for all analyzed values of *α* and *β.* Since *α*=0.07 and *β****=***-1 yielded the largest number of recognized cyclospectra in CYCLOLIBRARY (46 out of 81) and no cyclospectra in the LINEARLIBRARY and POLYMERLIBRARY datasets (Supplementary Figure S5), we selected these values as the default parameters.

**Supplementary Figure S5.**
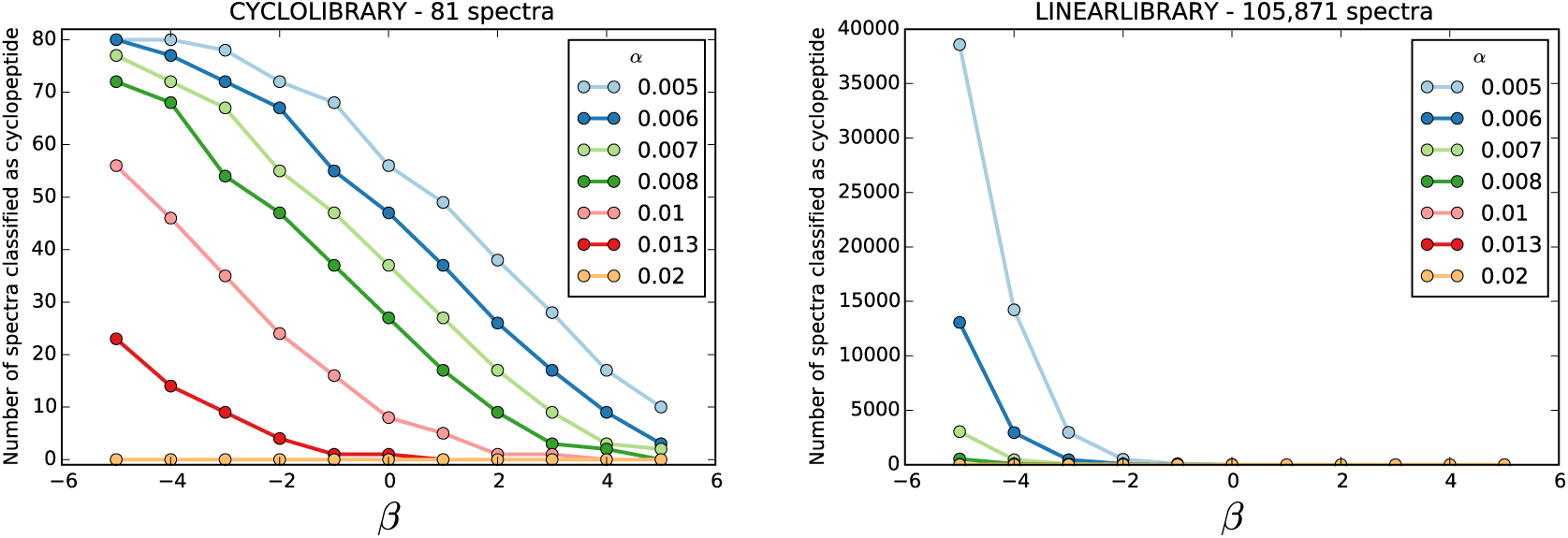
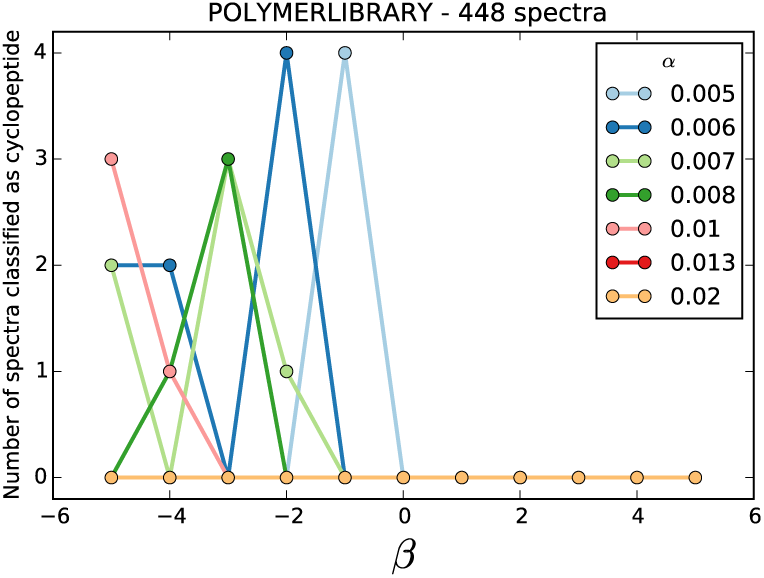
Number of spectra passing both the “high multiplicity cyclopeptidic cluster” and the “polymer” tests in the CYCLOLIBRARY, LINERALIBRARY, and POLYMERLiBRARY datasets (for various values of parameters *α* and *β*).

Supplementary Figure S6 presents the values of *cycloIntensity* and *k-merScore* for each spectrum in the CYCLOLIBRARY, LINEARLIBRARY, and POLYMERLIBRARY datasets and reveals a separation between the former and the two latter datasets with respect to these two parameters. CycloNovo thus classifies a spectrum as a cyclospectrum if its *cycloIntensity* exceeds the *cycloIntensity* threshold (60%) and its *k-merScore* exceeds the *k-merScore* threshold (5). 45 spectra in the CYCLOLIBRARY datasets, that pass all four tests described above, are classified as cyclospectra.

**Supplementary Figure S6.**
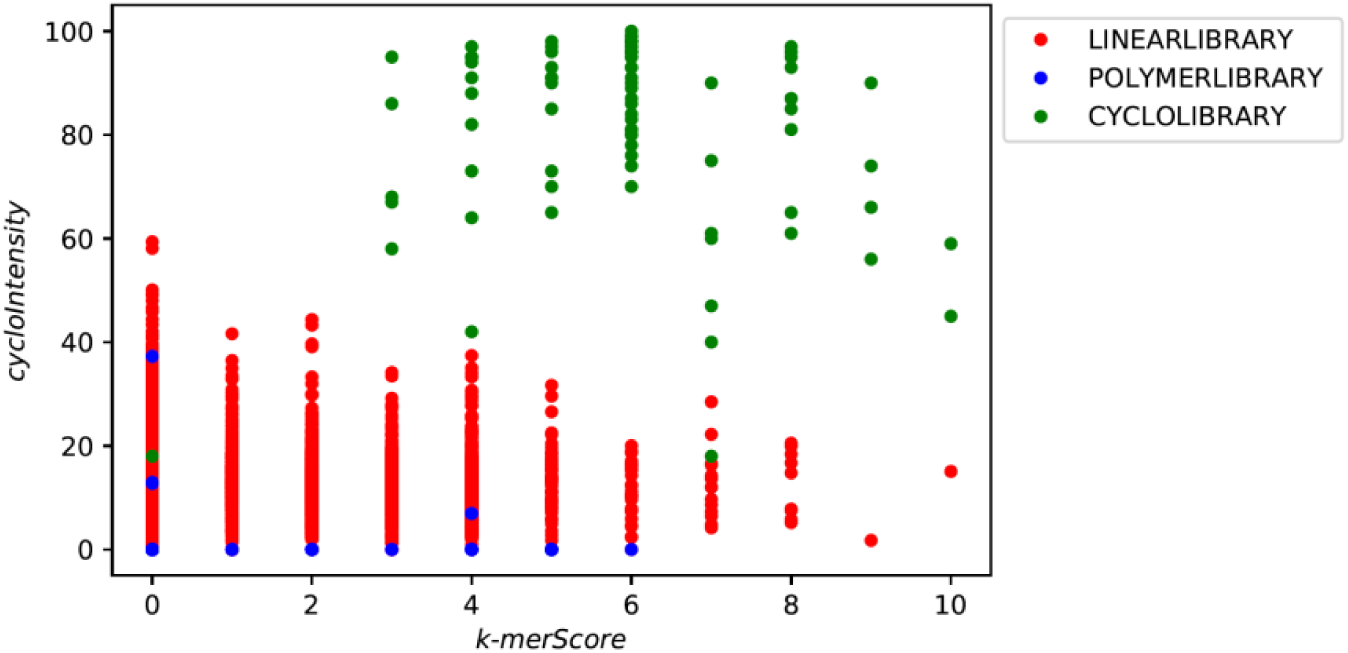
Values of *cycloIntensity* and *k-merScore* for all spectra in the CYCLOLIBRARY, POLYMERS, and LINEARLIBRARY datasets (for *k*=5).

We also investigated how CycloNovo’s ability to recognize a cyclospectrum is affected by the fragmentation quality of the corresponding PSM (measured by the P-value of this PSM). For each spectrum in the CYCLOLIBRARY dataset, we identified the minimum value of the parameter *α* that leads to classifying this spectrum as cyclopeptidic (for *β*=- 1). Supplementary Figure S7 illustrates that well-fragmented spectra can be recognized even with more restrictive threshold values (larger values of *α*).

**Supplementary Figure S7.**
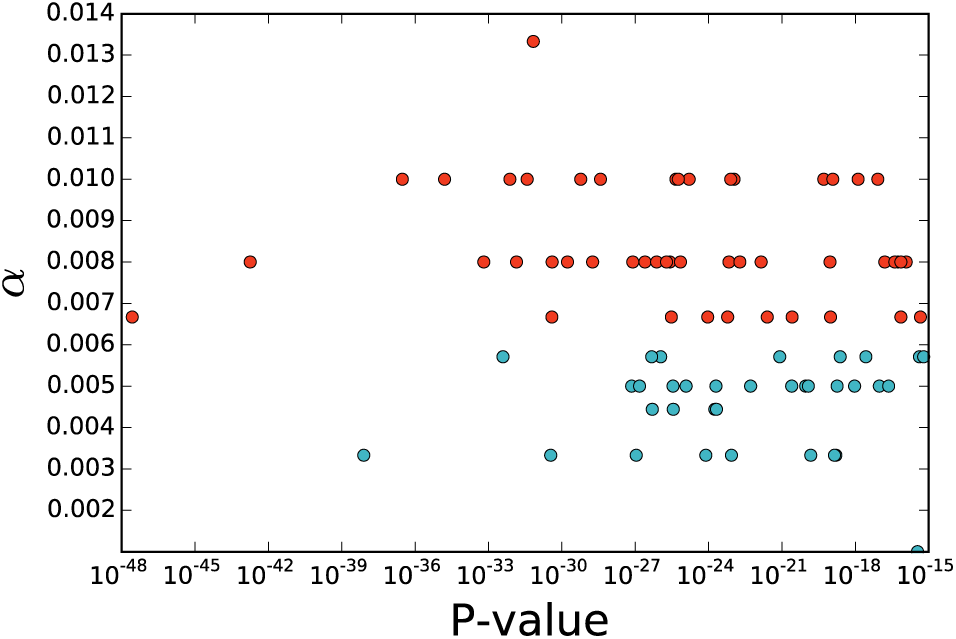
Dependence between the P-value of each spectrum in the CYCLOLIBRARY dataset and the minimum value of the parameter *α* that leads to classifying this spectrum as a cyclospectrum (for). Each point represents a spectrum in the CYCLOLIBRARY dataset. The *x*-axis shows the P-value of the PSM for that spectrum and the *y*-axis shows the minimum value of the parameter *α* that leads to classifying this spectrum as a cyclospectrum. The points corresponding to cyclospectra recognized with the default parameter *α*=0.07 are shown in red.

#### Supplementary Note: Cyclopeptidic amino acids

Table S2 lists all cyclopeptidic amino acids.

**Supplementary Table S2.** The list of 33 cyclopeptidic amino acids (corresponding to 31 unique amino acid masses). Proteinogenic amino acids are shown in blue, common amino acids in RiPPs^6^ are shown in black, and the remaining amino acids that appeared in the top 25 most frequent residues in *CyclopeptideDatabase* are shown in red.

| amino acid | elemental composition | monoisotopic mass (Da) | % of cyclopeptides in *CyclopeptideDatabase* containing the amino acid |
| --- | --- | --- | --- |
| isoleucine/leucine | C<sub>6</sub>H<sub>11</sub>ON | 113.084 | 55.8 |
| valine | C<sub>5</sub>H<sub>9</sub>ON | 99.068 | 38.2 |
| proline | C<sub>5</sub>H<sub>7</sub>ON | 97.053 | 36.9 |
| alanine | C<sub>3</sub>H<sub>5</sub>ON | 71.037 | 36.4 |
| phenylalanine | C<sub>9</sub>H<sub>9</sub>ON | 147.068 | 24.6 |
| methyl-isoleucine/leucine | C<sub>7</sub>H<sub>13</sub>ON | 127.101 | 18.7 |
| glycine | C<sub>2</sub>H<sub>3</sub>ON | 57.022 | 17 |
| threonine | C<sub>4</sub>H<sub>7</sub>O<sub>2</sub>N | 101.048 | 16.5 |
| serine | C<sub>3</sub>H<sub>5</sub>O<sub>2</sub>N | 87.032 | 15.8 |
| ornithine | C<sub>5</sub>H<sub>10</sub>ON<sup>2</sup> | 114.068 | 15 |
| methyl-alanine | C<sub>4</sub>H<sub>7</sub>ON | 85.054 | 14.6 |
| tyrosine | C<sub>9</sub>H<sub>9</sub>O<sub>2</sub>N | 163.063 | 10.9 |
| glutamine | C<sub>5</sub>H<sub>7</sub>O<sub>3</sub>N | 129.043 | 10.1 |
| asparagine | C<sub>4</sub>H<sub>6</sub>O<sub>2</sub>N<sub>2</sub> | 114.043 | 9.5 |
| methyl-phenylalanine | C<sub>10</sub>H<sub>11</sub>ON | 161.085 | 9.5 |
| aspartic acid | C<sub>4</sub>H<sub>5</sub>O<sub>3</sub>N | 115.027 | 7.6 |
| glutamic acid | C<sub>5</sub>H<sub>8</sub>O<sub>2</sub>N<sub>2</sub> | 128.059 | 7.6 |
| arginine | C<sub>6</sub>H<sub>12</sub>ON<sub>4</sub> | 156.101 | 7.2 |
| tryptophan | C<sub>11</sub>H<sub>10</sub>ON<sub>2</sub> | 186.079 | 7.2 |
| methyl-oxazoline | C<sub>4</sub>H<sub>7</sub>NO | 85.104 | 2.9 |
| lysine | C<sub>6</sub>H<sub>12</sub>ON<sub>2</sub> | 128.095 | 2.9 |
| oxazoline | C<sub>3</sub>H<sub>5</sub>NO | 71.078 | 2.5 |
| methionine | C<sub>5</sub>H<sub>9</sub>ONS | 131.040 | 2.4 |
| oxazole | C<sub>3</sub>H<sub>3</sub>NO | 69.062 | 1.3 |
| methionine-oxide | C<sub>5</sub>H<sub>9</sub>O<sub>2</sub>NS | 147.040 | 1.2 |
| histidine | C<sub>6</sub>H<sub>7</sub>ON<sub>3</sub> | 137.059 | 1 |
| methionine-dioxide | C<sub>5</sub>H<sub>9</sub>O<sub>3</sub>NS | 163.040 | 0.3 |
| methyl-oxazole | C<sub>4</sub>H<sub>5</sub>NO | 83.089 | <0.1 |
| thiazole | C<sub>3</sub>H<sub>3</sub>NS | 85.128 | <0.1 |
| thiazoline | C<sub>3</sub>H<sub>5</sub>NS | 87.143 | <0.1 |
| cysteine | C<sub>3</sub>H<sub>5</sub>ONS | 103.009 | <0.1 |

#### Supplementary Note: Analyzing spectral convolution

Supplementary Figure S8 illustrates that each amino acid in surugamide results in a populous diagonal in the spectral diagram of *Spectrum*_*Surugamide*_. For each constructed cluster (diagonal band in the spectral diagram), we consider all pairs of masses in *Spectrum* that contributed to this cluster and form a *band* as the set of these *k* pairs.

We define the *cluster diameter* as the difference between its maximum and minimum elements. Supplementary Figure S9 presents the band for the cluster with multiplicity 8 and mass 128.09 (diameter 0.03) in the spectral convolution of *Spectrum*_*Surugamide*_and reveals that the 8 elements of this band can be partitioned into 7 groups of closely located points. We are interested in the number of such groups (rather than the raw cluster multiplicities) since experimental spectra often contain *satellite masses* resulting from *neutral losses* and *isotopic peaks.* For example, in addition to the integer mass 242 Da corresponding to the peptide IK, *Spectrum*_*Surugamide*_also contains the integer mass 225 Da corresponding to the loss of NH_3_from this peptide.

|  |  |  |  |  |
| --- | --- | --- | --- | --- |
| $y \approx x + 128$ | (185.13, 313.22)<br>**(446.30, 574.38)** | (261.17, 389.26)<br>**(445.28, 573.35)** | (298.21, 426.28)<br>(487.35, 615.42) | (374.24, 502.32)<br>(784.52, 912.62) |

**Supplementary Figure S8.**
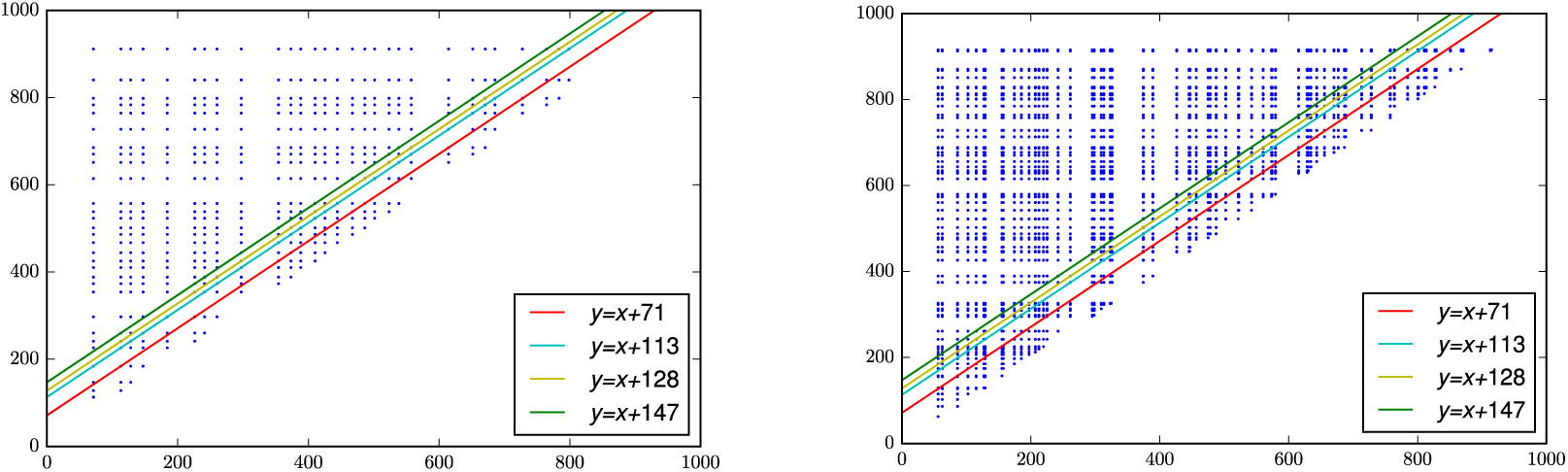
The spectral diagrams of the *TheoreticalSpectrum(Surugamide)* (left) and *SpectrumSurugamide* (right). The highlighted lines with slope +1 have *y*-intercepts equal to the masses of the constituent amino acids of surugamide (A, L/I, K, and F). Amino acids A, L/I, K, and F correspond to populous diagonals containing 11, 23, 11, and 11 points (left figure) and 5, 14, 8, and 4 points (right figure), respectively.

**Supplementary Figure S9.**
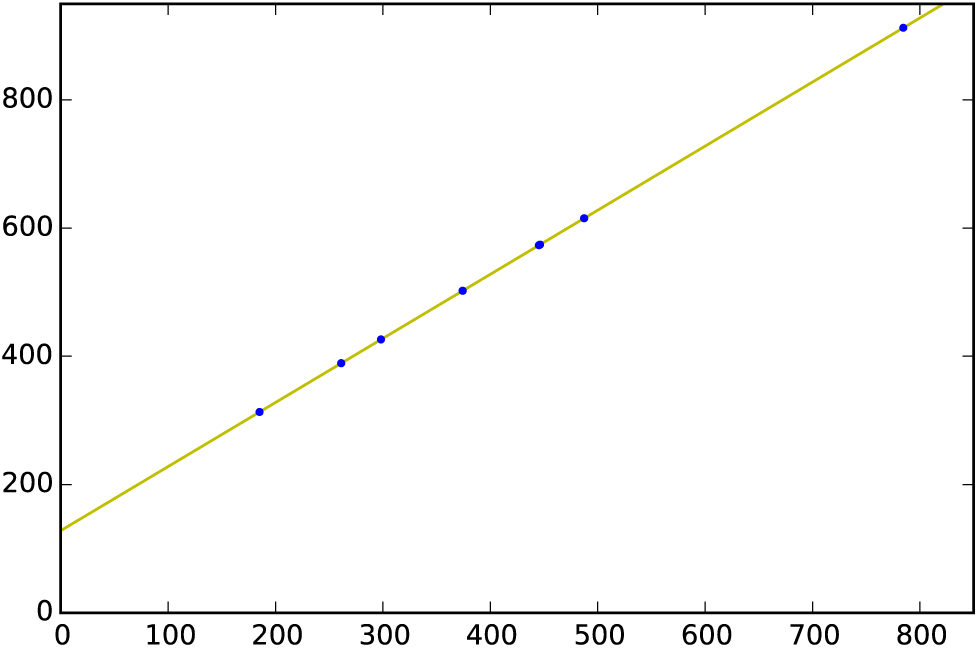
A band with multiplicity eight in *SpectrumSurugamide* (cluster with mass 128.09 and diameter 0.03). (Top) Coordinates of the points in the band. Since the difference between the *x*-coordinates and *y*-coordinates of the two points shown in bold match the mass of hydrogen, these two points are clustered together in this band. (Bottom) The same band in the spectral diagram for *Spectrum*_*Surugamide*_. The points of the band can be partitioned into seven groups of closely located points: six singleton groups and one group with two elements. H)NHCO3

Since satellite masses artificially inflate cluster multiplicities, there is a need to reduce biases caused by these masses. We thus define the set of common satellite offsets (1 Da (, 18 Da (H_2_O), 17 Da (), and 28 Da ()) and perform additional single linkage clustering in each populous band by combining pairs of masses in a single cluster if both their *x*-coordinates and *y*-coordinates differ by a satellite offset. We redefine the concept of cluster multiplicity as the number of the resulting clusters in the band (Supplementary Table S3).

**Supplementary Table S3.** List of clusters in the spectral convolution of *SpectrumSurugamide*. Clusters are shown in the decreasing order of their multiplicities (only clusters with multiplicity at least 2 are shown). Cyclopeptidic clusters are shown in bold and cyclopeptidic clusters with masses similar to masses of amino acids in surugamide are shown in red.

| cluster mass | multiplicity after satellite removal | multiplicity before satellite removal | cluster diameter | cluster distance |
| --- | --- | --- | --- | --- |
| **113.078** | **12** | **14** | **0.106** | **0.006 (I/L)** |
| **128.089** | **7** | **8** | **0.032** | **0.006 (K)** |
| 56.025 | 6 | 6 | 0.033 | - |
| **186.108** | **6** | **6** | **0.077** | **0.029 (W)** |
| **71.041** | **5** | **5** | **0.022** | **0.004 (A)** |
| **147.069** | **4** | **4** | **0.027** | **0.001 (F)** |
| **156.099** | **4** | **5** | **0.036** | **0.002 (R)** |
| 184.106 | 4 | 5 | 0.032 | - |
| 98.079 | 4 | 4 | 0.031 | - |
| 169.11 | 4 | 5 | 0.01 | - |
| 57.049 | 4 | 4 | 0.02 | - |
| 70.042 | 3 | 3 | 0.01 | - |
| 132.058 | 3 | 3 | 0.006 | - |
| 133.584 | 3 | 3 | 0.013 | - |
| 183.116 | 3 | 3 | 0.024 | - |
| 168.159 | 3 | 3 | 0.023 | - |
| 52.049 | 3 | 3 | 0.005 | - |
| 96.035 | 3 | 3 | 0.017 | - |
| 165.115 | 3 | 3 | 0.004 | - |
| 76.041 | 3 | 4 | 0.013 | - |
| 73.08 | 3 | 3 | 0.017 | - |
| **101.053** | **3** | **3** | **0.011** | **0.005 (T)** |
| 167.886 | 3 | 4 | 0.024 | - |
| 189.105 | 2 | 3 | 0.009 | - |
| **57.018** | **2** | **2** | **0.006** | **0.004 (G)** |
| 114.09 | 2 | 2 | 0.01 | - |
| 45.041 | 2 | 2 | 0.012 | - |

#### Supplementary Note: De Bruijn graphs for *Spectrum*_*Surugamide*_

Supplementary Figure S10 shows the pruned *de Bruijn* graphs of three compositions of *Spectrum*_*Surugamide*_that do not contain feasible cycles.

**Supplementary Figure S10.**
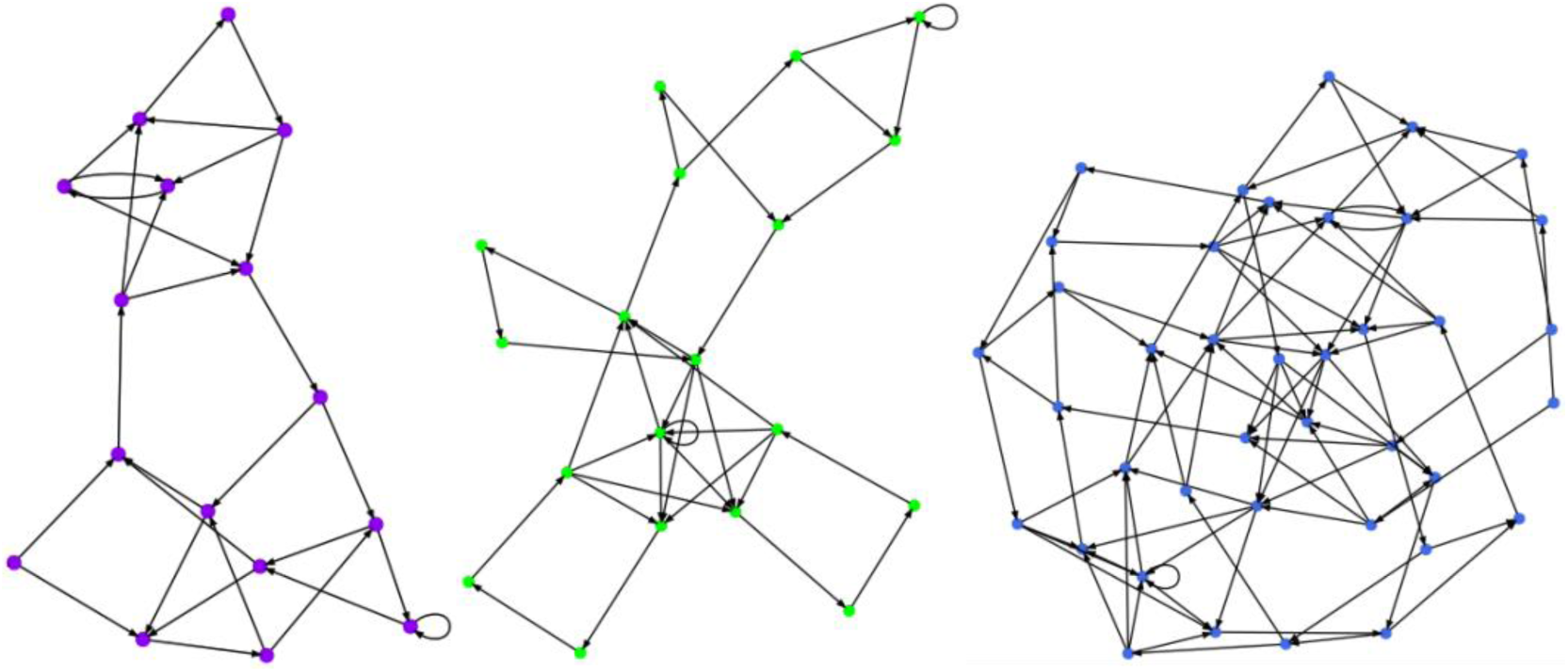
The pruned *de Bruijn* graphs of the compositions of *Spectrum*. _*Surugamide*_ **that do not contain feasible cycles**. (Left) The composition 113^4^156^2^147^1^ results in a de Bruijn graph with 40 vertices and 57 edges and a pruned de Bruijn graph with 18 vertices and 40 edges. (Middle) The composition 128^4^71^2^156^1^101^1^ results in a de Bruijn graph with 52 vertices and 76 edges and a pruned de Bruijn graph with 20 vertices and 42 edges. (Right) The composition 128^5^113^1^101^1^57^1^ results in a de Bruijn graph with 94 vertices and 180 edges and a pruned de Bruijn graph with 40 vertices and 92 edges.

#### Supplementary Note: CycloNovo running time

CycloNovo recognizes cyclospectra by constructing their spectral convolutions (*O*(*n*^*2*^) running time for a spectrum with *n* peaks) and further sequences all found cyclospectra. Using a single 2.5GHz processor, CycloNovo recognized all cyclospectra in the HUMANSTOOL and GNPS datasets in ≈35 minutes and ≈31 hours, respectively.

The sequencing step is only applied to a small fraction of all spectra in spectral datasets, e.g., CycloNovo recognizes only ≈0.05% of all spectra in the GNPS dataset as cyclospectra. The running time for the sequencing step varies widely between spectra and depends on the number of putative amino acid compositions, the number of putative *k*-mers, and the number feasible cycles in the de Bruijn graphs.

We were not able to benchmark CycloNovo against CYCLONE^7^ since CYCLONE failed to reconstruct most spectra in the CYCLOLIBRARY dataset. For example, CycloNovo took ≈3 seconds to sequence *Spectrum*^*Surugamide*^. In contrast, CYCLONE^7^ failed to sequence *Spectrum*^*Surugamide*^ and was not even able to infer alanine and lysine as amino acids in suragamide.

We thus compared CycloNovo with a brute force sequencing algorithm by generating and scoring all possible permutations of all amino acid compositions resulting from the putative amino acids for *Spectrum*_*Surugamide*_. The brute-force approach took ≈27 seconds to sequence *Spectrum*_*Surugamide*_as the highest scoring reconstruction among all 113 generated cyclopeptides. In a more difficult example, we analyzed the spectrum with precursor mass 899.36 and reconstructed the orbitide FVDTTGYD in the S.VULGARIS dataset (Table 2). CycloNovo sequenced this spectrum in 56 seconds while the brute force approach took 58 minutes. For this spectrum, there exist 321 putative amino acid compositions yielding over 400,000 candidate sequences. Since the majority of those ∼400,000 sequences do not produce high-scoring 5-mers, they yield the relatively small de Bruijn graphs and hence small numbers of feasible cycles and candidate cyclopeptides. By only exploring the feasible cycles in the de Bruijn graphs, CycloNovo reduced the number of candidate sequences to 2,491.

These examples illustrate that the running time of CycloNovo varies by orders of magnitude depending on analyzed cyclospectra. By only exploring the sequences spelled by the feasible cycles in the de Bruijn graphs, CycloNovo greatly reduces the search space compared to the brute force approach. For example, the brute force approach failed after 1000 hours on just two cyclospectra from the CYCLOLIBRARY dataset, while the de Bruijn graph approach finished analysis of all spectra in this library in ≈48 hours. Supplementary Note “CycloNovo analysis of the CYCLOLIBRARY dataset” lists all CycloNovo reconstructions for this dataset. CycloNovo analysis of the HUMANSTOOL and GNPS datasets took ∼34 and ∼149 hours, respectively.

#### Supplementary Note: Information about spectral datasets

##### Information about *CyclopeptideDatabase*

The *PNPDatabase*^8^ combines all known peptidic natural products from various databases. Many peptides in this database are *lipopeptides* containing a lipid chain, e.g., surfactin is a cyclopeptide containing a fatty acid side chain connected to a fully peptidic part via a peptide bond.

We classify a peptide in the *PNPdatabase* as a cyclopeptide if its backbone could be represented as a circular graph (cycle) with nodes corresponding to either a single amino acid or a single lipid tail (i.e. monomers) and edges corresponding to the amide bonds in the peptide structure. 1,257 out of 5,021 peptides in the *PNPDatabase* represent cyclopeptides and form *CyclopeptideDatabase* (note that the *CyclopeptideDatabase* database contains lipopeptides).

##### Information about the CYCLOLIBRARY dataset

We searched ∼130 million GNPS spectra against the *CyclopeptideDatabase* using Dereplicator^1^ and identified 81 distinct cyclopeptides (41 cyclofamilies) corresponding to PSMs with FDR=0% and P-value below 10^−15^. For each identified cyclopeptide, we selected the PSM with the minimum P-value (among all PSMs identified for this cyclopeptide), resulting in a set of 81 PSMs and hence created a spectral dataset CYCLOLIBRARY with 81 spectra (Table S4). CYCLOLIBRARY includes only 13 cyclopeptides (6 cyclofamilies) that are made up entirely of cyclopeptidic amino acids (Table S4). 34 peptides (25 cyclofamilies) in the CYCLOLIBRARY dataset contain lipid tails and 34 peptides (14 cyclofamilies) contain non-cyclopeptidic amino acids.

**Supplementary Table S4.**
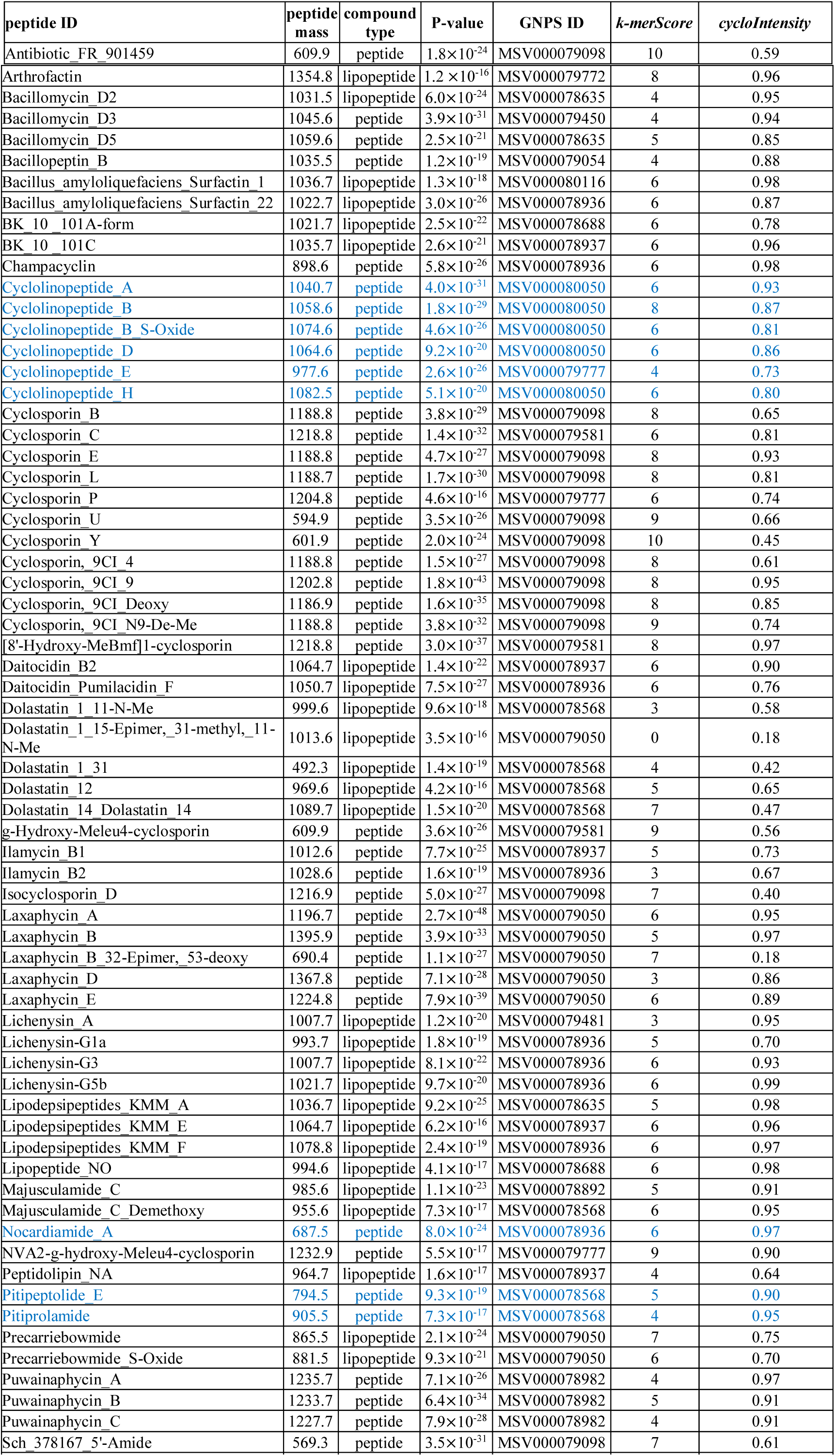
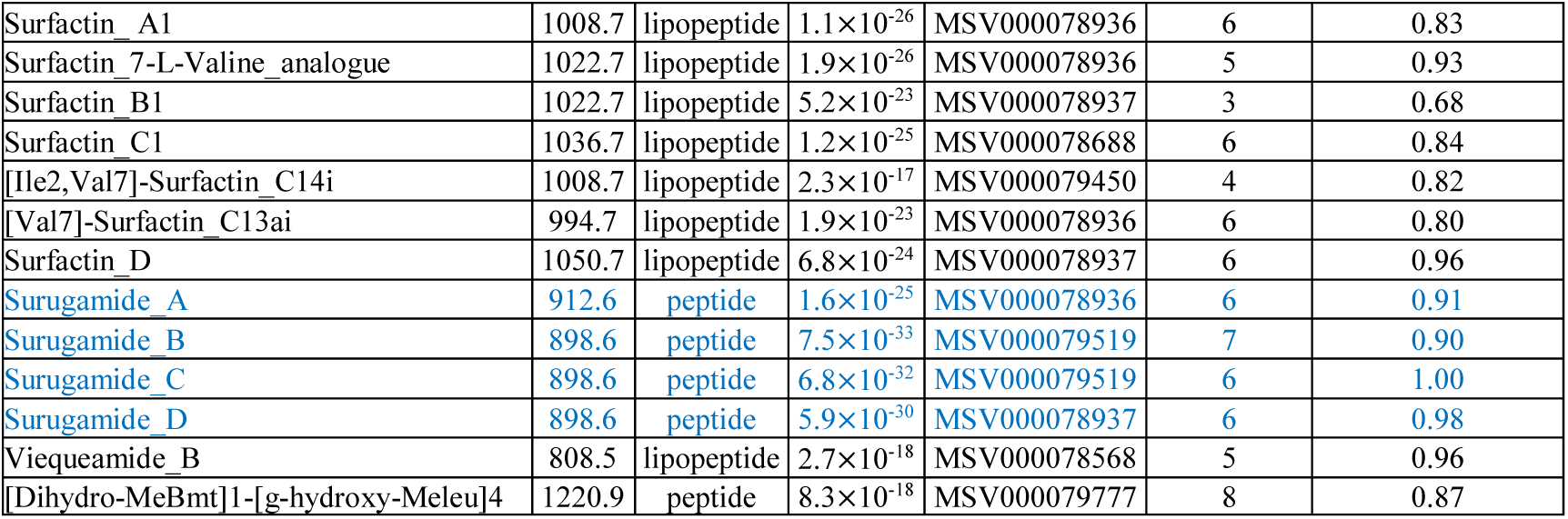
Cyclopeptides in the CYCLOLIBRARY dataset. The peptides that gave rise to 81 spectra in the CYCLOLIBRARY dataset with their corresponding peptide mass, P-value, the GNPS ID of the dataset a spectrum belongs to, *k-merScore*, and *cycloIntensity*. The column “compound type” specifies whether the compound is fully peptidic or represents a lipopeptide. The blue rows show the 13 cyclopeptides that are made up entirely of cyclopeptidic amino acids.

##### Information about the GNPS dataset

The GNPS dataset is formed by 40 MassIVE datasets that were selected from 120 datasets analyzed in Gurevich et al.^8^ to exclude potentially miscalibrated spectral datasets. Since miscalibrated datasets typically do not result in any cyclopeptide identifications, we searched each of these 120 datasets with Dereplicator and excluded datasets that did not result in any identifications (with 0% FDR and P-value below 10^−15^) from further analysis, leaving us with 40 datasets (Supplementary Table S5).

**Supplementary Table S5.** Information about the GNPS dataset. The last row shows the total number of spectra and unique cyclopeptides/cyclofamiles across all datasets. The datasets marked in red, blue, and green form GNPS^CYANO^, GNPS_PSEUDO_, and GNPS_ACTI_ subsets of the GNPS dataset, respectively.

| GNPS ID | #spectra | #spectra after pre-processing | #cyclo spectra | #putative cyclo-peptides/ cyclo-families found by CycloNovo | #identified cyclo-peptides/ cyclo-families identified by Dereplicator (among cyclo-spectra) | #identified cyclo-peptides/ cyclo-families identified by Dereplicator (among all spectra) | #identified branch-cyclic peptides/ branch-cyclic families identified by Dereplicator (among cyclo-spectra) |
| --- | --- | --- | --- | --- | --- | --- | --- |
| MSV000078567 | 730582 | 316993 | 4 | 2/1 | 2/1 | 4/2 | 0/0 |
| MSV000078568 | 23582408 | 12118472 | 317 | 74/35 | 9/8 | 15/10 | 1/1 |
| MSV000078584 | 680906 | 263160 | 0 | 0/0 | 0/0 | 0/0 | 0/0 |
| MSV000078604 | 311617 | 281617 | 606 | 56/25 | 6/3 | 6/3 | 3/3 |
| MSV000078606 | 289170 | 237988 | 122 | 32/12 | 1/1 | 1/1 | 7/6 |
| MSV000078635 | 680168 | 569316 | 2388 | 124/40 | 9/4 | 12/5 | 10/7 |
| MSV000078656 | 2844 | 1023 | 88 | 14/1 | 10/5 | 11/6 | 3/3 |
| MSV000078710 | 1469076 | 689912 | 6 | 1/1 | 2/2 | 3/3 | 0/0 |
| MSV000078787 | 1767830 | 1281235 | 208 | 58/31 | 25/13 | 25/13 | 8/7 |
| MSV000078839 | 717600 | 504350 | 1 | 1/1 | 1/1 | 1/1 | 0/0 |
| MSV000078847 | 167917 | 115603 | 19 | 7/5 | 1/1 | 1/1 | 1/1 |
| MSV000078892 | 847114 | 461769 | 27 | 8/4 | 3/2 | 4/3 | 0/0 |
| MSV000078936 | 2059306 | 1538683 | 526 | 58/30 | 25/13 | 30/15 | 5/5 |
| MSV000078937 | 1694918 | 1303349 | 256 | 52/26 | 26/14 | 33/19 | 11/9 |
| MSV000078982 | 984 | 727 | 32 | 4/2 | 3/1 | 3/1 | 2/2 |
| MSV000079044 | 576282 | 270860 | 2 | 1/1 | 1/1 | 2/1 | 0/0 |
| MSV000079050 | 1241328 | 683124 | 207 | 24/8 | 3/1 | 7/3 | 3/3 |
| MSV000079054 | 702020 | 364382 | 112 | 16/7 | 13/6 | 13/6 | 3/3 |
| MSV000079069 | 847145 | 215229 | 1066 | 23/2 | 0/0 | 1/1 | 0/0 |
| MSV000079140 | 607488 | 443147 | 1118 | 25/7 | 14/6 | 15/7 | 0/0 |
| MSV000079274 | 5433248 | 3457806 | 14 | 2/2 | 1/1 | 1/1 | 0/0 |
| MSV000079312 | 54806 | 25354 | 1112 | 15/3 | 0/0 | 1/1 | 0/0 |
| MSV000079450 | 697812 | 581012 | 2245 | 120/39 | 6/2 | 6/2 | 9/6 |
| MSV000079471 | 22379 | 16138 | 68 | 14/4 | 10/4 | 10/4 | 0/0 |
| MSV000079481 | 45742 | 7692 | 48 | 2/1 | 0/0 | 2/2 | 0/0 |
| MSV000079502 | 47450 | 3167 | 14 | 3/1 | 3/1 | 4/2 | 0/0 |
| MSV000079516 | 120154 | 19113 | 138 | 22/6 | 4/2 | 5/2 | 0/0 |
| MSV000079517 | 22516 | 2911 | 37 | 7/3 | 3/1 | 4/2 | 0/0 |
| MSV000079519 | 76289 | 13985 | 112 | 15/6 | 3/1 | 5/2 | 0/0 |
| MSV000079568 | 224645 | 10273 | 0 | 0/0 | 0/0 | 4/2 | 0/0 |
| MSV000079581 | 129012 | 41779 | 74 | 4/3 | 2/2 | 5/5 | 0/0 |
| MSV000079598 | 919494 | 286130 | 9 | 3/1 | 4/1 | 6/3 | 0/0 |
| MSV000079651 | 81818 | 5239 | 0 | 0/0 | 0/0 | 2/1 | 0/0 |
| MSV000079679 | 595244 | 300682 | 109 | 24/14 | 12/7 | 14/7 | 2/2 |
| MSV000079772 | 75916 | 13870 | 40 | 7/5 | 2/2 | 2/2 | 0/0 |
| MSV000079778 | 1242178 | 451962 | 265 | 50/44 | 6/1 | 7/2 | 0/0 |
| MSV000079813 | 578683 | 170990 | 23 | 6/4 | 2/2 | 3/2 | 0/0 |
| MSV000079888 | 238820 | 74317 | 170 | 17/7 | 8/4 | 9/5 | 1/1 |
| MSV000080115 | 1567520 | 709527 | 400 | 33/9 | 12/6 | 13/7 | 9/6 |
| MSV000080116 | 70250 | 31009 | 19 | 9/5 | 6/3 | 6/3 | 0/0 |
| **TOTAL** | 51220679 | 27883895 | 12004 | 512/213 | 61/37 | 91/51 | 41/27 |

##### Supplementary Note: CycloNovo analysis of the CYCLOLIBRARY dataset

CycloNovo recognized 45 out of 81 spectra in the CYCLOLIBRARY dataset as cyclospectra. It classified 12 out of 13 cyclopeptides built from cyclopeptidic amino acids as cyclospectra and *de novo* sequenced them with one of the top three highest scores.

CycloNovo is unable to sequence most spectra in the CYCLOLIBRARY dataset since 68 of them originated from lipopeptides or peptides containing non-cyclopeptidic amino acids. To evaluate how CycloNovo performs on 45 cyclospectra in this dataset, we extended the set of cyclopeptidic amino acids to include the mass of the lipid chain and/or the masses of non-cyclopeptidic amino acids for each spectrum. Using this admittedly imperfect benchmarking approach, CycloNovo sequenced 22 of 45 cyclospectra as a highest-scoring *de novo* reconstructions and an additional 16 spectra with one of the three highest scores. Supplementary Table S6 lists the highest-scoring reconstruction for these spectra and illustrates that the highest-scoring reconstruction is similar to the correct amino acid sequence for all these spectra.

**Supplementary Table S6.**
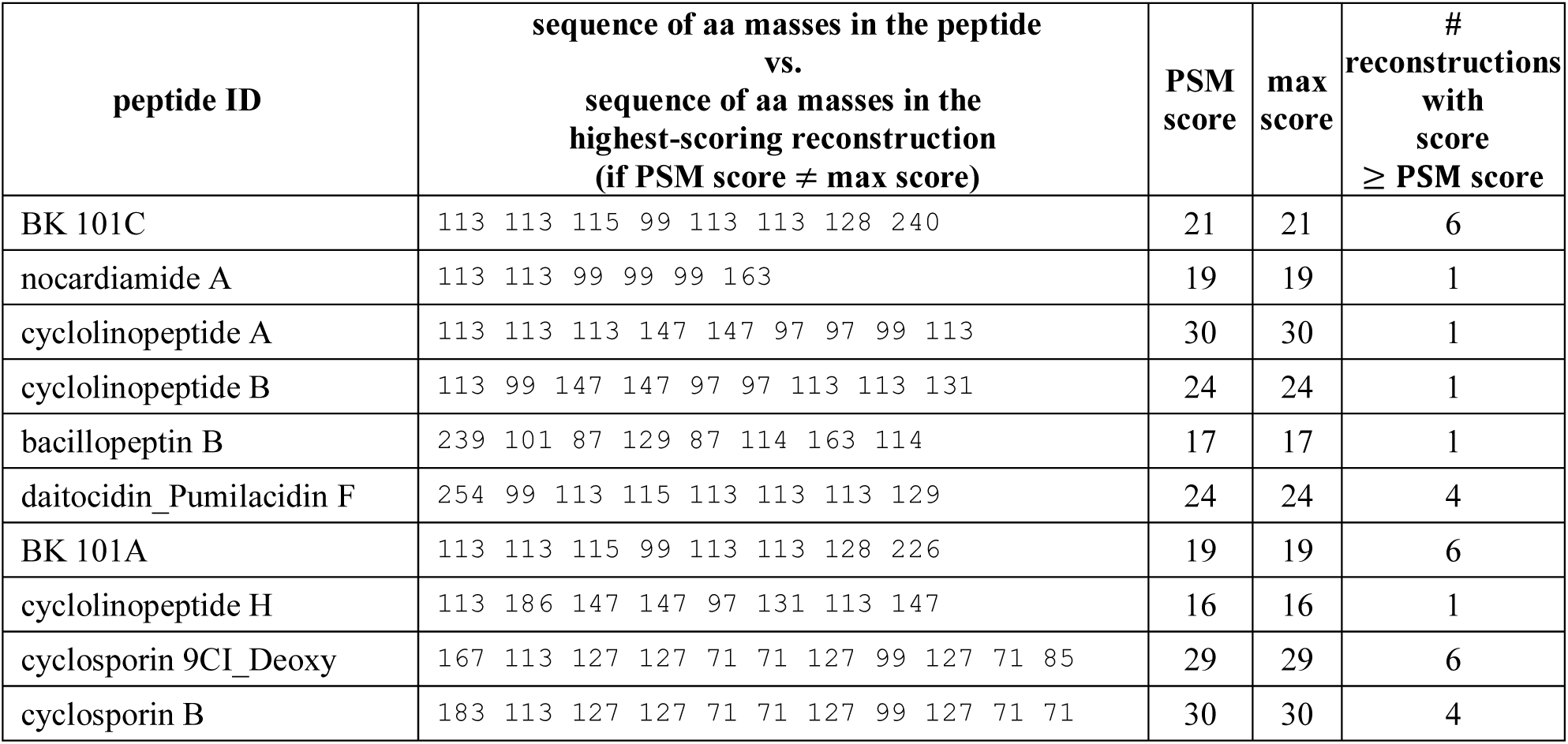
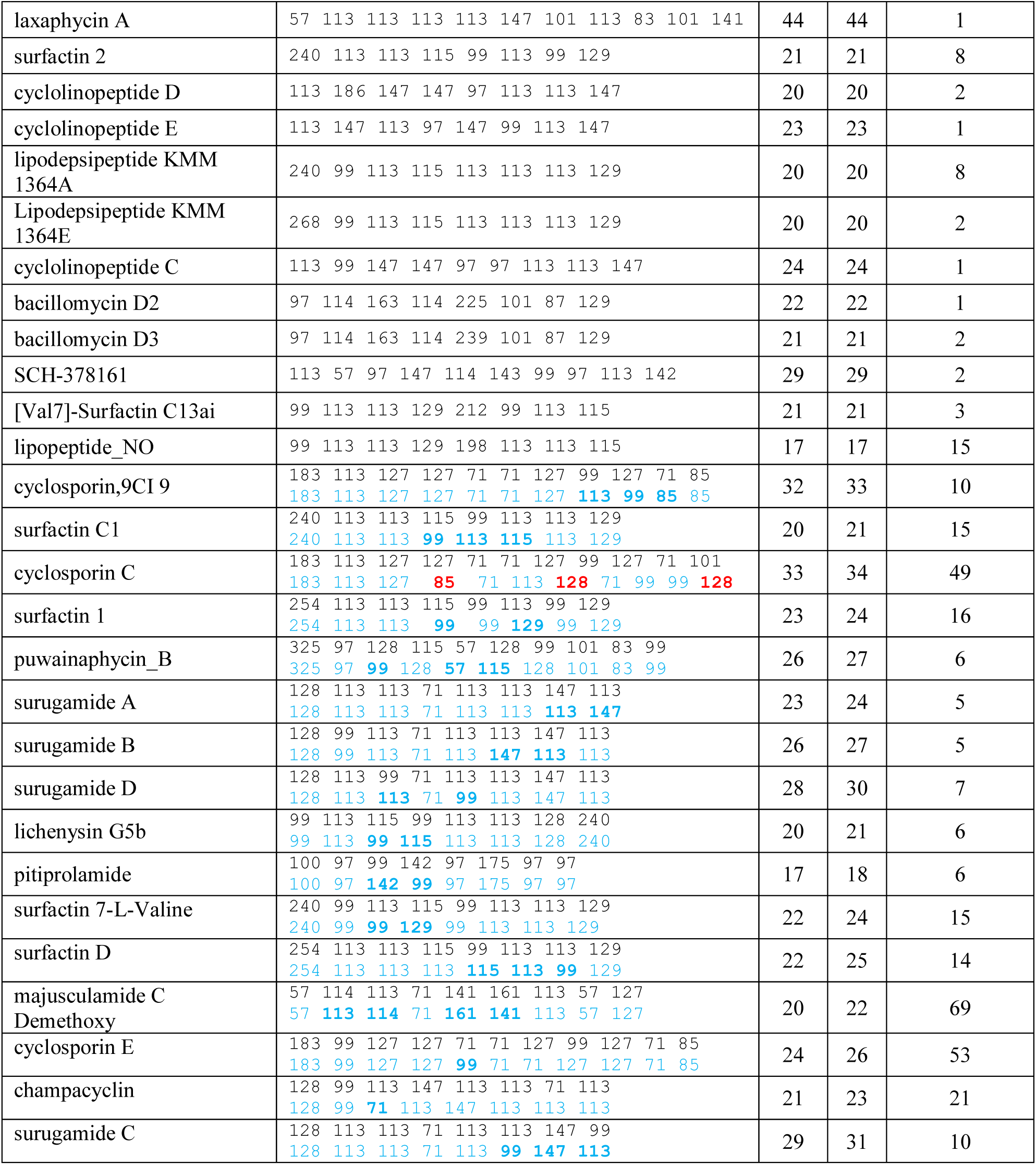
38 cyclopeptides reconstructed by CycloNovo from 45 cyclospectra in the CYCLOLIBRARY dataset. The PSM score represents the score of the PSM in the CYCLOLIBRARY dataset. The “max score” represents the score of the top-scoring reconstruction. For 22 cyclopeptides, the correct sequence of the cyclopeptide has the highest-scoring reconstruction. For the remaining 16 cyclopeptides, a highest-scoring reconstruction is listed below the correct sequence of the cyclopeptide in blue (differently arranged amino acid masses in the reconstructed cyclopeptide are shown in bold blue). Only in one case (cyclosporin C), CycloNovo predicted the wrong amino acids (shown in red) for the top-scoring reconstruction. CycloNovo failed to sequences 45-38=7 cyclospectra in the CYCLOLIBRARY dataset since it was not able to predict all their amino acids.

##### Supplementary Note: Cyclopeptide-encoding transcripts in the S.VULGARIS dataset

Supplementary Table S7 lists ORFs (translated into amino acid sequences) in the orbitide-encoding transcripts. The PawL1 proteins have dual fates; they encode an albumin as well as a cyclopeptide(s). An enzyme asparaginyl endopeptidase (targets Asp, Asn) matures both the albumin and the cyclopeptide.

**Supplementary Table S7.**
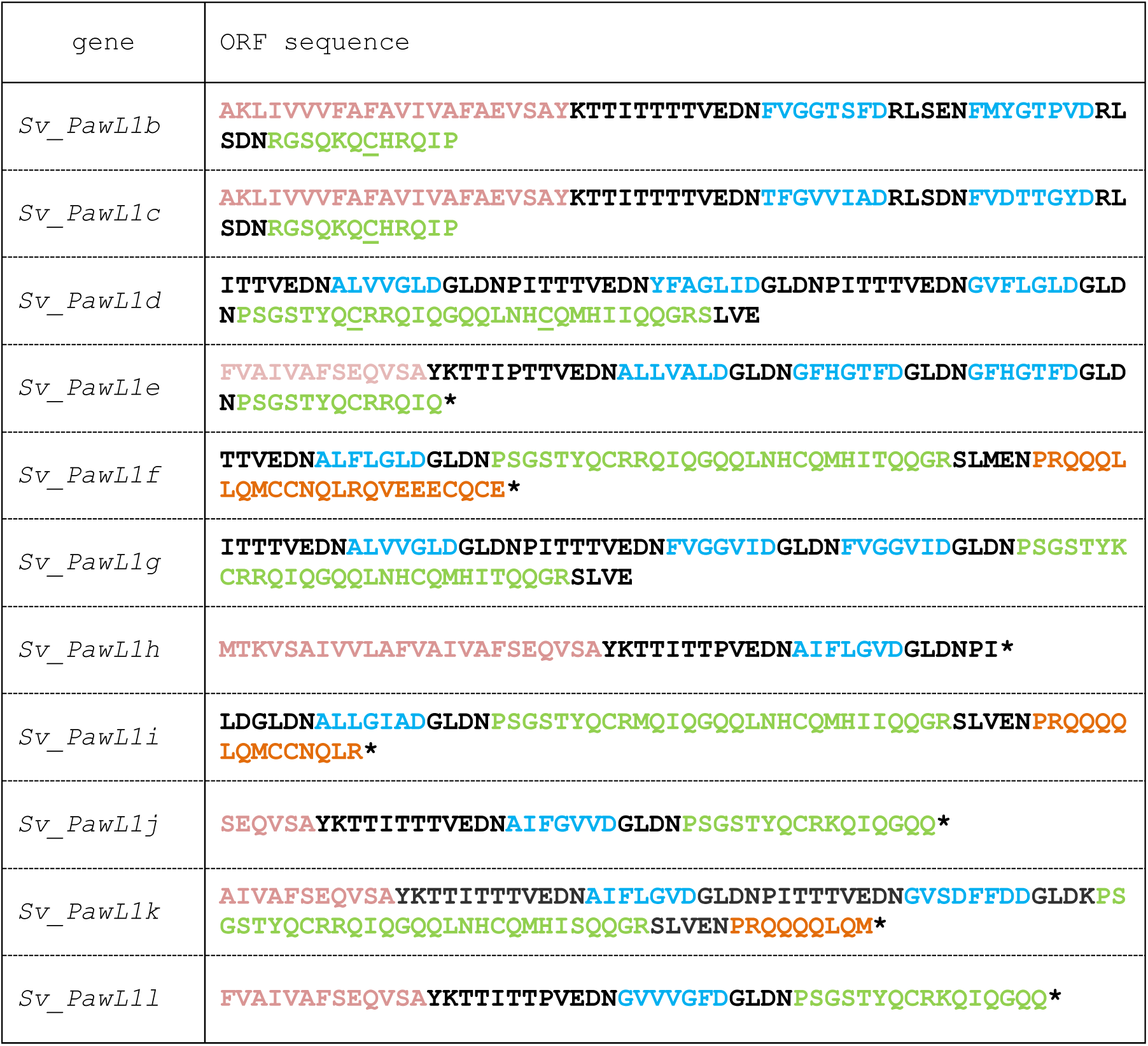
ORFs in the cyclopeptide-encoding transcripts. All identified ORFs originate from various *PawS1-Like* genes. The sequences are color-coded based on the subunits they belong to: endoplasmic reticulum signal sequence (pink), the reconstructed cyclopeptide (blue), 2S albumin small subunit (lime green), and 2S albumin large subunit (orange). While the first three sequences (*Sv_PawL1b, Sv_PawL1c*, and *Sv_PawL1d*) are known *PawS1-Like* genes in *S. vulgaris*, the other eight sequences (named *Sv_PawL1e* through *Sv_ PawL1l*) are novel *PawS1-Like* genes that were identified by searching for novel cyclopeptides.

**Supplementary Figure S11.**
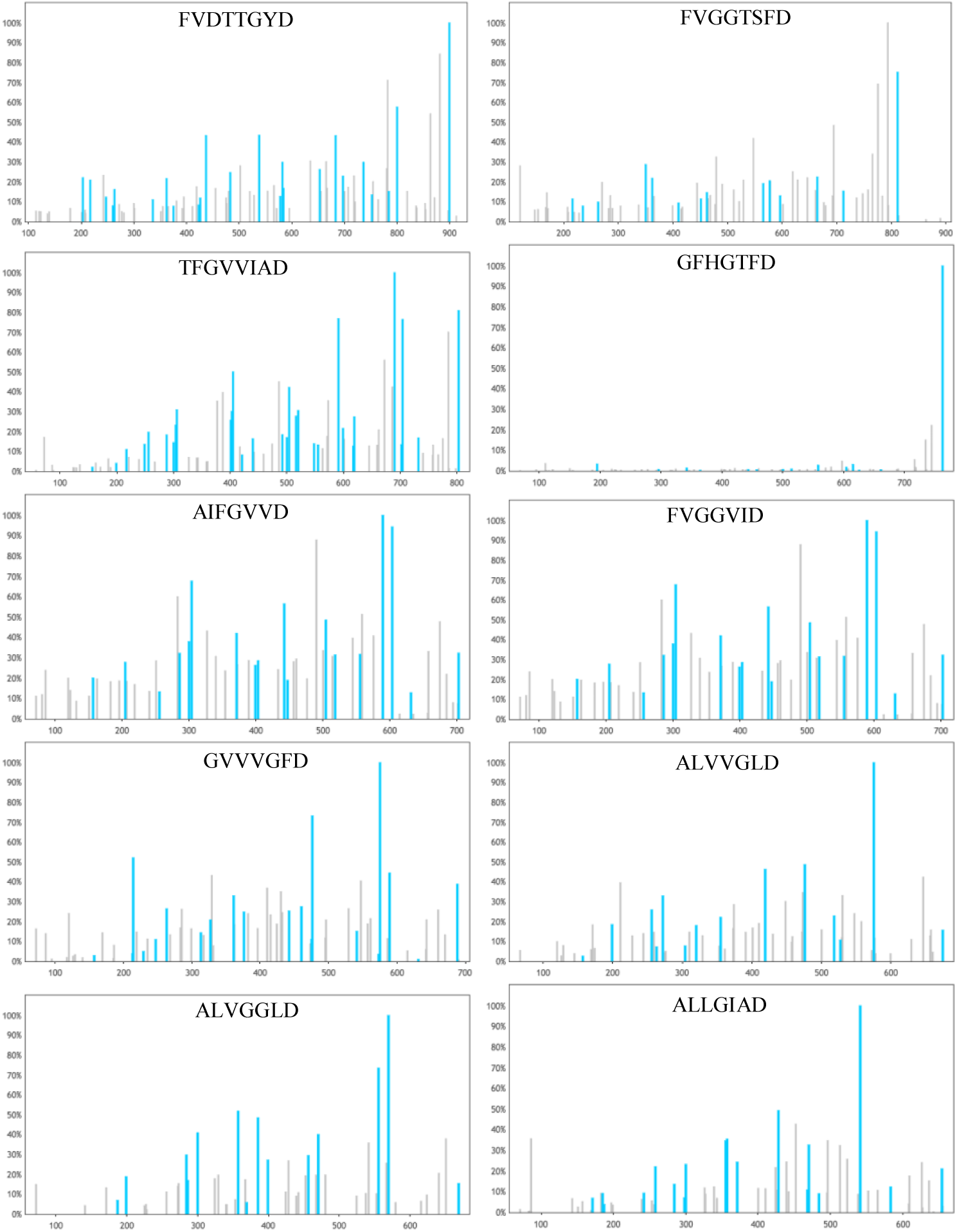
Annotated cyclospectra of the ten reconstructed cyclopeptides in the S.VULGARIS dataset. The *x*-axis shows the *m/z* ratios and the *y*-axis shows the percentage of the peak intensity compared to the intensity of the largest peak in that spectrum.

Supplementary Figure S11 shows cyclospectra in the S.VULGARIS dataset, annotated using their CycloNovo reconstructions.

###### Identification (Dereplicator) and *de novo* reconstruction (CycloNovo) of peptides in the HUMANSTOOL dataset

Supplementary Table S8 lists cyclopeptides identified by Dereplicator in the HUMANSTOOL datasets. Supplementary Table S9 lists CycloNovo reconstructions of 31 cyclopeptides in the HUMANSTOOL dataset.

**Supplementary Table S8.** Cyclopeptides identified by Dereplicator in the HUMANSTOOL dataset. The correct sequence of all reconstructed cyclopeptides has the highest score among all reconstructions. For each cyclopeptide, the score of the correct cyclopeptide (column “PSM score”), the number of reconstructions with scores larger or equal to the PSM score (column “#reconstructions score score”), and P-values are listed.

| precursor mass | peptide | PSM score | #reconstructions with score ≥ PSM score | P-value | peptide ID |
| --- | --- | --- | --- | --- | --- |
| 1040.66 | ILVPPFFLI | 31 | 1 | $1.2 \times 10^{-54}$ | cyclolinopeptide A |
| 1058.61 | MLIPPFVI | 24 | 1 | $2.3 \times 10^{-42}$ | cyclolinopeptide B |
| 1074.62 | M<sup>+16</sup>LIPPFVI | 16 | 1 | $9.7 \times 10^{-18}$ | cyclolinopeptide C |
| 1064.57 | M<sup>+16</sup>LLPFFWI | 20 | 2 | $1.5 \times 10^{-33}$ | cyclolinopeptide D |
| 1082.52 | M<sup>+16</sup>LMPFFWI | 19 | 1 | $1.2 \times 10^{-31}$ | cyclolinopeptide H |
| 977.56 | M<sup>+16</sup>LVFPLFI | 25 | 1 | 1.6×10<sup>-43</sup> | cyclolinopeptide E |
| 961.55 | MLVFPLFI | 25 | 10 | 3.1×10<sup>-42</sup> | cyclolinopeptide P |
| 567.36 | GIVIPS | 11 | 1 | 2.1×10<sup>-17</sup> | citrusin V |

**Supplementary Table S9.**
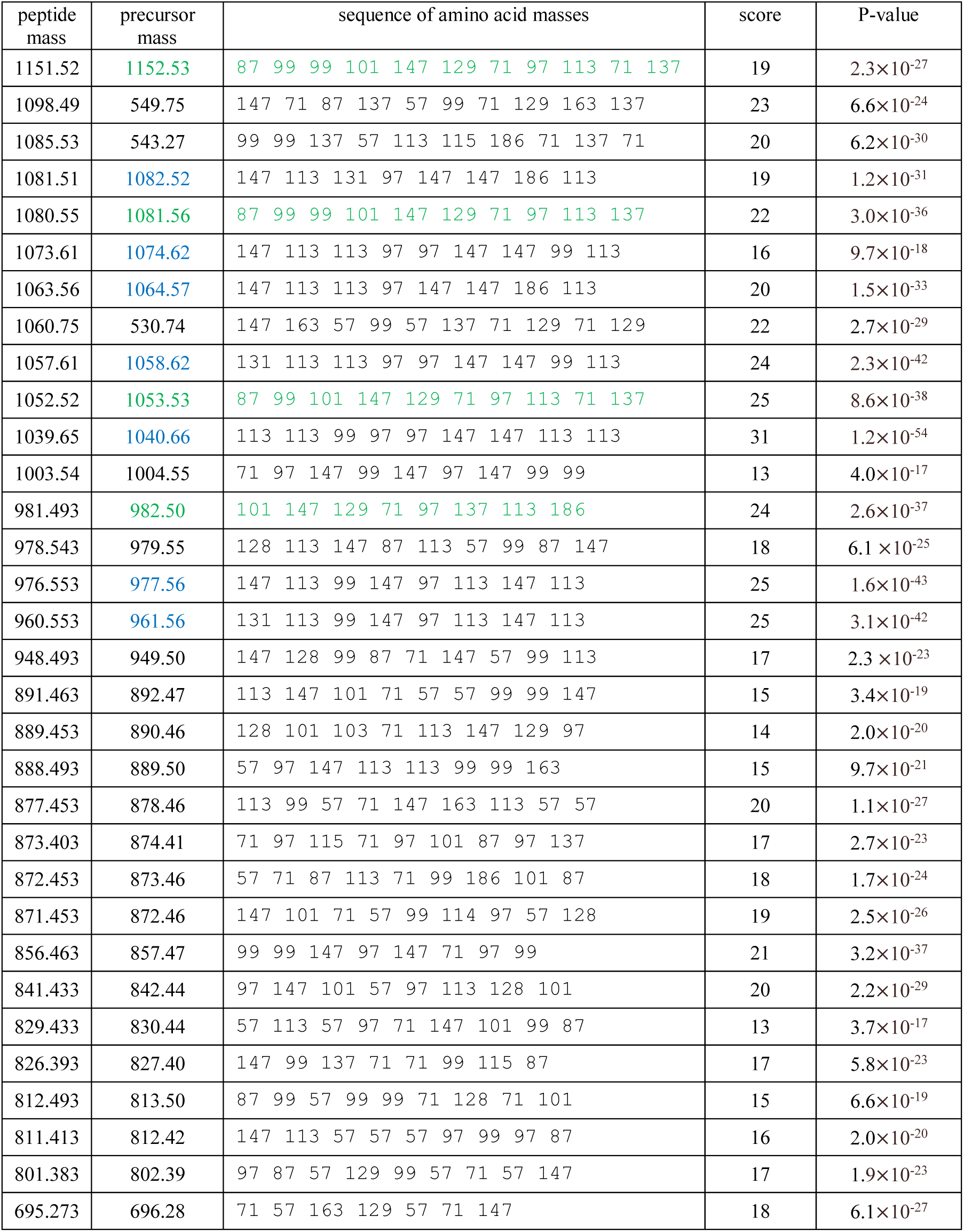
*De novo* reconstructions of 31 cyclopeptides in the HUMANSTOOL dataset. For each spectrum, its precursor mass, the *de novo* reconstruction (shown as a sequence of nominal masses of amino acids), the score, and the P-value are shown. *De novo* reconstructions are ordered in the decreasing order of their precursor masses. Precursor masses of spectra identified by Dereplicator are highlighted in blue. Cyclopeptides highlighted in green represent a novel cyclofamily described in the Supplementary Note “Novel cyclofamily in the HUMANSTOOL dataset.”

| peptide mass | precursor mass | sequence of amino acid masses | score | P-value |
| --- | --- | --- | --- | --- |
| 1151.52 | 1152.53 | 87 99 99 101 147 129 71 97 113 71 137 | 19 | 2.3×10<sup>-27</sup> |
| 1098.49 | 549.75 | 147 71 87 137 57 99 71 129 163 137 | 23 | 6.6×10<sup>-24</sup> |
| 1085.53 | 543.27 | 99 99 137 57 113 115 186 71 137 71 | 20 | 6.2×10<sup>-30</sup> |
| 1081.51 | 1082.52 | 147 113 131 97 147 147 186 113 | 19 | 1.2×10<sup>-31</sup> |
| 1080.55 | 1081.56 | 87 99 99 101 147 129 71 97 113 137 | 22 | 3.0×10<sup>-36</sup> |
| 1073.61 | 1074.62 | 147 113 113 97 97 147 147 99 113 | 16 | 9.7×10<sup>-18</sup> |
| 1063.56 | 1064.57 | 147 113 113 97 147 147 186 113 | 20 | 1.5×10<sup>-33</sup> |
| 1060.75 | 530.74 | 147 163 57 99 57 137 71 129 71 129 | 22 | 2.7×10<sup>-29</sup> |
| 1057.61 | 1058.62 | 131 113 113 97 97 147 147 99 113 | 24 | 2.3×10<sup>-42</sup> |
| 1052.52 | 1053.53 | 87 99 101 147 129 71 97 113 71 137 | 25 | 8.6×10<sup>-38</sup> |
| 1039.65 | 1040.66 | 113 113 99 97 97 147 147 113 113 | 31 | 1.2×10<sup>-54</sup> |
| 1003.54 | 1004.55 | 71 97 147 99 147 97 147 99 99 | 13 | 4.0×10<sup>-17</sup> |
| 981.493 | 982.50 | 101 147 129 71 97 137 113 186 | 24 | 2.6×10<sup>-37</sup> |
| 978.543 | 979.55 | 128 113 147 87 113 57 99 87 147 | 18 | 6.1×10<sup>-25</sup> |
| 976.553 | 977.56 | 147 113 99 147 97 113 147 113 | 25 | 1.6×10<sup>-43</sup> |
| 960.553 | 961.56 | 131 113 99 147 97 113 147 113 | 25 | 3.1×10<sup>-42</sup> |
| 948.493 | 949.50 | 147 128 99 87 71 147 57 99 113 | 17 | 2.3×10<sup>-23</sup> |
| 891.463 | 892.47 | 113 147 101 71 57 57 99 99 147 | 15 | 3.4×10<sup>-19</sup> |
| 889.453 | 890.46 | 128 101 103 71 113 147 129 97 | 14 | 2.0×10<sup>-20</sup> |
| 888.493 | 889.50 | 57 97 147 113 113 99 99 163 | 15 | 9.7×10<sup>-21</sup> |
| 877.453 | 878.46 | 113 99 57 71 147 163 113 57 57 | 20 | 1.1×10<sup>-27</sup> |
| 873.403 | 874.41 | 71 97 115 71 97 101 87 97 137 | 17 | 2.7×10<sup>-23</sup> |
| 872.453 | 873.46 | 57 71 87 113 71 99 186 101 87 | 18 | 1.7×10<sup>-24</sup> |
| 871.453 | 872.46 | 147 101 71 57 99 114 97 57 128 | 19 | 2.5×10<sup>-26</sup> |
| 856.463 | 857.47 | 99 99 147 97 147 71 97 99 | 21 | 3.2×10<sup>-37</sup> |
| 841.433 | 842.44 | 97 147 101 57 97 113 128 101 | 20 | 2.2×10<sup>-29</sup> |
| 829.433 | 830.44 | 57 113 57 97 71 147 101 99 87 | 13 | 3.7×10<sup>-17</sup> |
| 826.393 | 827.40 | 147 99 137 71 71 99 115 87 | 17 | 5.8×10<sup>-23</sup> |
| 812.493 | 813.50 | 87 99 57 99 99 71 128 71 101 | 15 | 6.6×10<sup>-19</sup> |
| 811.413 | 812.42 | 147 113 57 57 57 97 99 97 87 | 16 | 2.0×10<sup>-20</sup> |
| 801.383 | 802.39 | 97 87 57 129 99 57 71 57 147 | 17 | 1.9×10<sup>-23</sup> |
| 695.273 | 696.28 | 71 57 163 129 57 71 147 | 18 | 6.1×10<sup>-27</sup> |

###### Cyclospectra of branch-cyclic peptides in the HUMANSTOOL dataset

We classify a peptide as branch-cyclic if its backbone includes a cycle (with all monomers connected via amide bonds) and a side chain that includes at least one additional amide bond not included in the cycle. Although CycloNovo classify spectra of some branch cyclic peptides as cyclospectra (see Supplementary Note “Information about spectral datasets”), it is unable to *de novo* sequence them. Nevertheless, CycloNovo provides information about substrings of branch-cyclic peptides made of cyclopeptidic amino acids. For example, CycloNovo classified the spectrum of massetolide F in the HUMANSTOOL dataset as a cyclospectrum. The lipopetide massetolide F consists of the cycle TILSLSLV and a branch EL (along with a fatty acid chain tail with nominal mass 171 Da) connected to the cycle via an amide bond between T and E. We represent this branch cyclic peptides as a concatenate between the sequence of nominal masses of the cyclic and branch region separated by “*” sign, i.e., massetolide F is represented as 100, 113, 87, 113, 87, 113, 99 * 129, 113, 171. CycloNovo found five cyclopeptidic amino acids in massetolide F (S, I, L, V, T, and E) and missed the lipid chain (171 Da).

###### Assembly of the human stool sample where massetolide F was detected

We used metaSPAdes^10^ to assemble the metagenomic dataset, generated from the stool sample (dated by 6/16/2014) where massetolide F was detected, This dataset includes 34.5 million paired reads which are assembled into 81 thousand scaffolds of lengths longer than 500 bp amounting to 407 Mb total assembly length.

###### Supplementary Note: Cyclopeptides in the GNPS dataset

Supplementary Figure S13 shows the number of identified cyclopeptides across all GNPS sub-datasets. Supplementary Figure S14 shows the number of cyclopeptides and cyclofamilies that gave rise to cyclospectra found by CycloNovo across all GNPS sub-datasets.

**Supplementary Figure S13.**
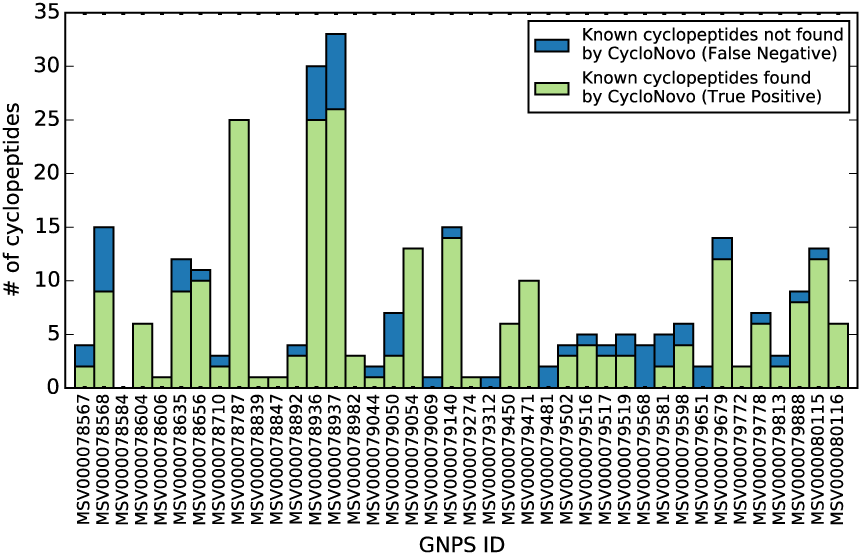
Number of cyclopeptides identified by Dereplicator across all GNPS sub-dataset. Dereplicator identified 81 cyclopeptides in the GNPS dataset. Since some cyclopeptides are identified in multiple sub-datasets, the total numbers of identified cyclopeptides across all GNPS sub-datasets (180) exceeds 81. The green (blue) part of each bar represent spectra that were (were not) classified by CycloNovo as cyclospectra.

**Supplementary Figure S14.**
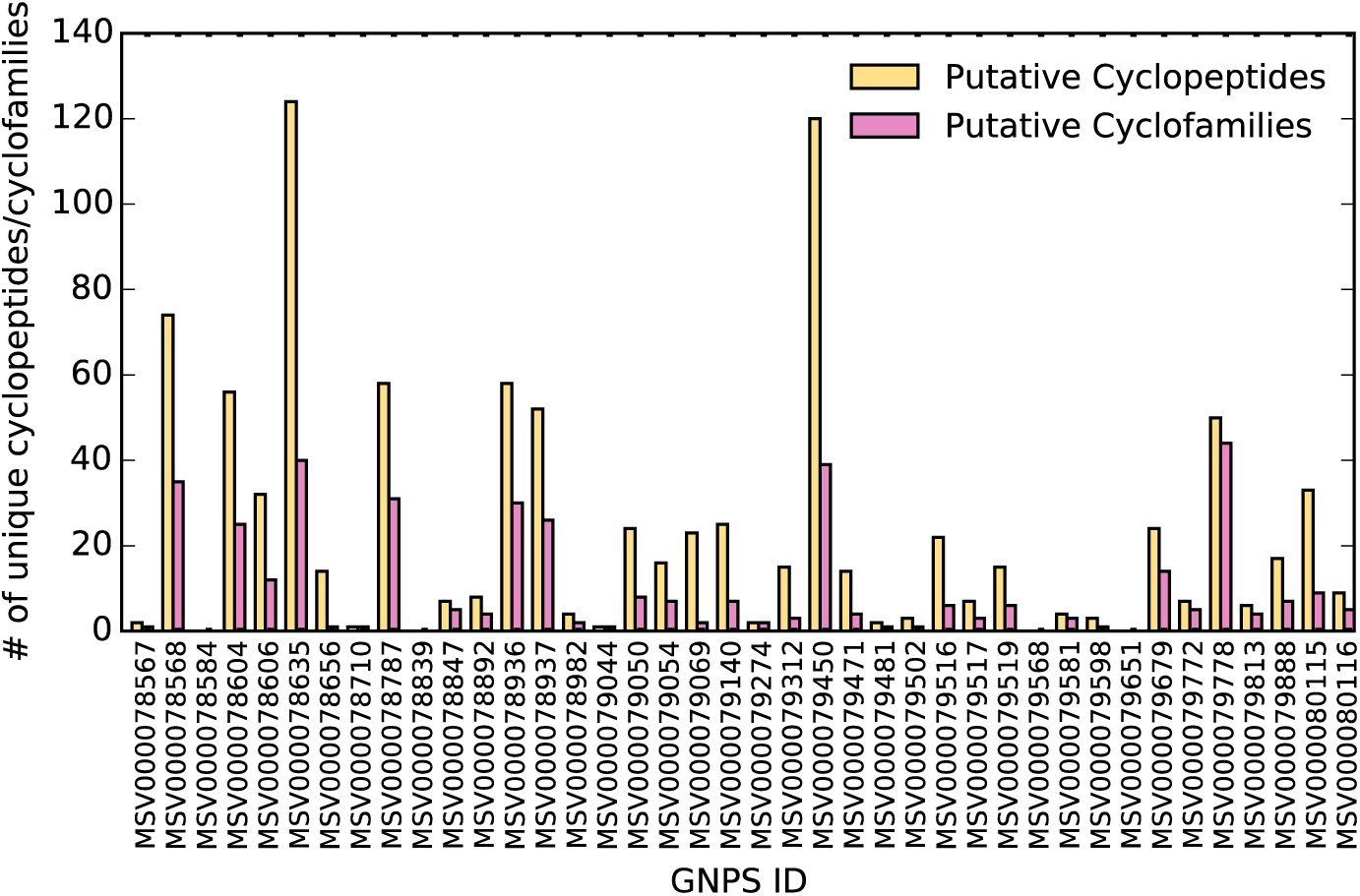
Number of cyclopeptides (yellow) and cyclofamilies (pink) found by CycloNovo across all GNPS dataset.

## REFERENCES

1. Ling, L. L. et al. A new antibiotic kills pathogens without detectable resistance. Nature 517, 455–459 (2015).

2. Medema, M. H. & Fischbach, M. A. Computational approaches to natural product discovery. Nat. Chem. Biol. 11, 639–648 (2015).

3. Wang, M. et al. Sharing and community curation of mass spectrometry data with Global Natural Products Social Molecular Networking. Nat. Biotechnol. 34, 828–837 (2016).

4. Mohimani, H. et al. Dereplication of peptidic natural products through database search of mass spectra. Nat. Chem. Biol. 13, 30–37 (2017).

5. Gurevich, A. et al. Increased diversity of peptidic natural products revealed by modification-tolerant database search of mass spectra. Nat. Microbiol. 3, 319–327 (2018).

6. Marahiel, M. A., Stachelhaus, T. & Mootz, H. D. Modular Peptide Synthetases Involved in Nonribosomal Peptide Synthesis. Chem. Rev. 97, 2651–2674 (1997).

7. Mohimani, H. et al. NRPquest: coupling mass spectrometry and genome mining for nonribosomal peptide discovery. J. Nat. Prod. 77, 1902–1909 (2014).

8. Mohimani, H. et al. Automated genome mining of ribosomal peptide natural products. ACS Chem. Biol. 9, 1545–1551 (2014).

9. Bandeira, N., Tsur, D., Frank, A. & Pevzner, P. A. Protein identification by spectral networks analysis. Proc. Natl. Acad. Sci. U.S.A. 104, 6140–6145 (2007).

10. Mohimani, H., Wei-Ting Liu, Y.-L. Y., Susana P. Gaudêncio, W. F., Dorrestein, P. C. & Pevzner, P. A. Multiplex de novo sequencing of peptide antibiotics. J. Comput. Biol. 18, 1371–1381 (2011).

11. Mohimani, H. & Pevzner, P. A. Dereplication, sequencing and identification of peptidic natural products: from genome mining to peptidogenomics to spectral networks. Nat. Prod. Rep. 33, 73–86 (2016).

12. Mohimani, H. et al. Cycloquest: Identification of cyclopeptides via database search of their mass spectra against genome databases. J. Proteome Res. 10, 4505–4512 (2011).

13. Ibrahim, A. et al. Dereplicating nonribosomal peptides using an informatic search algorithm for natural products (iSNAP) discovery. Proc. Natl. Acad. Sci. U.S.A. 109, 19196–19201 (2012).

14. Mohimani, H. et al. Sequencing cyclic peptides by multistage mass spectrometry. Proteomics 11, 3642–3650 (2011).

15. Ng, J. et al. Dereplication and de novo sequencing of nonribosomal peptides. Nat Methods 6, 596–599 (2009).

16. Kavan, D., Kuzma, M., Lemr, K., Schug, K. A. & Havlicek, V. CYCLONE - A utility for de novo sequencing of microbial cyclic peptides. Journal of the American Society for Mass Spectrometry 24, 1177–1184 (2013).

17. Townsend, C. et al. CycLS: Accurate, whole-librarysequencing of cyclic peptides using tandem mass spectrometry. Bioorg. Med. Chem. 26, 1232–1238 (2018).

18. Compeau, P. E. C., Pevzner, P. A. & Tesler, G. How to apply de Bruijn graphs to genome assembly. Nat. Biotechnol. 29, 987–991 (2011).

19. Takada, K. et al. Surugamides A − E, Cyclic Octapeptides with Four D-Amino Acid Residues, from a Marine Streptomyces sp.: LC-MS-Aided Inspection of Partial Hydrolysates for the Distinction of D- and L-Amino Acid Residues in the Sequence. 50, 3–7 (2013).

20. Bhushan, R. & Bruckner, H. Use of Marfey’s reagent and analogs for chiral amino acid analysis: Assessment and applications to natural products and biological systems. Journal of Chromatography B: Analytical Technologies in the Biomedical and Life Sciences 879, 3148–3161 (2011).

21. Mohimani, H., Kim, S. & Pevzner, P. A. A new approach to evaluating statistical significance of spectral identifications. Journal of Proteome Research 12, 1560–1568 (2013).

22. Jayasena, A. S. et al. Stepwise Evolution of a Buried Inhibitor Peptide over 45 My. Mol. Biol. Evol. 34, 1505–1516 (2017).

23. Bankevich, A. et al. SPAdes: A New Genome Assembly Algorithm and Its Applications to Single-Cell Sequencing. J. Comput. Biol. 19, 455–477 (2012).

24. Fisher, M. F. et al. A family of small, cyclic peptides buried in preproalbumin since the Eocene epoch. Plant Direct 2, e00042 (2018).

25. Mylne, J. S. et al. Albumins and their processing machinery are hijacked for cyclic peptides in sunflower. Nat. Chem. Biol. 7, 257–259 (2011).

26. Elliott, A. G. et al. Evolutionary Origins of a Bioactive Peptide Buried within Preproalbumin. Plant Cell 26, 981–995 (2014).

27. Yazdani, M. et al. Using machine learning to identify major shifts in human gut microbiome protein family abundance in disease. in Proceedings - 2016 IEEE International Conference on Big Data, Big Data 2016 1272–1280 (2016).

28. Frank, A. M. et al. Spectral archives: extending spectral libraries to analyze both identified and unidentified spectra. Nat. Methods 8, 587–591 (2011).

29. Noh, H. J. et al. Anti-inflammatory activity of a new cyclic peptide, citrusin XI, isolated from the fruits of Citrus unshiu. J. Ethnopharmacol. 163, 106–112 (2015).

30. Belknap, W. R. et al. A family of small cyclic amphipathic peptides (SCAmpPs) genes in citrus. BMC Genomics 16, 303 (2015).

31. Kaufmann, H. P. & Tobschirbel, A. Über ein oligopeptid aus leinsamen. Eur. J. Inorg. Chem. 92, 2805–2809 (1959).

32. Morita, H. et al. A new immunosuppressive cyclic nonapeptide, cycloinopeptide B from Linum usitatissimum. Bioorganic Med. Chem. Lett. 7, 1269–1272 (1997).

33. Morita, H., Shishido, A., Matsumoto, T., Itokawa, H. & Takeya, K. Cyclolinopeptides B-E, new cyclic peptides from Linum usitatissimum. Tetrahedron 55, 967–976 (1999).

34. Okinyo-Owiti, D. P., Young, L., Burnett, P. G. G. & Reaney, M. J. T. New flaxseed orbitides: Detection, sequencing, and15N incorporation. Biopolym. - Pept. Sci. Sect. 102, 168–175 (2014).

35. Gerard, J. et al. Massetolides A-H, antimycobacterial cyclic depsipeptides produced by two pseudomonads isolated from marine habitats. J. Nat. Prod. 60, 223–229 (1997).

36. Scales, B. S., Dickson, R. P., Lipuma, J. J. & Huffnagle, G. B. Microbiology, genomics, and clinical significance of the Pseudomonas fluorescens species complex, an unappreciated colonizer of humans. Clin. Microbiol. Rev. 27, 927–948 (2014).

37. O’Sullivan, D. J. & O’Gara, F. Traits of fluorescent Pseudomonas spp. involved in suppression of plant root pathogens. Microbiol. Rev. 56, 662–676 (1992).

38. Watrous, J. et al. Mass spectral molecular networking of living microbial colonies. Proc. Natl. Acad. Sci. U.S.A. 109, E1743–E1752 (2012).

39. Nguyen, D. D. et al. Indexing the Pseudomonas specialized metabolome enabled the discovery of poaeamide B and the bananamides. Nat. Microbiol. 2, (2016).

40. Charlop-Powers, Z. et al. Urban park soil microbiomes are a rich reservoir of natural product biosynthetic diversity. Proc. Natl. Acad. Sci. (2016).

41. Nothias, L.-F., Knight, R. & Dorrestein, P. C. Antibiotic discovery is a walk in the park. Proc. Natl. Acad. Sci. (2016).

42. Bommareddy, A. et al. Effects of dietary flaxseed on intestinal tumorigenesis in Apc Min mouse. Nutr. Cancer 61, 276–283 (2009).

43. Shim, Y. Y., Gui, B., Arnison, P. G., Wang, Y. & Reaney, M. J. T. Flaxseed (Linum usitatissimum L.) bioactive compounds and peptide nomenclature: Areview. Trends in Food Science and Technology 38, 5–20 (2014).

44. Adolphe, J. L., Whiting, S. J., Juurlink, B. H. J., Thorpe, L. U. & Alcorn, J. Health effects with consumption of the flax lignan secoisolariciresinol diglucoside. Br. J. Nutr. 103, 929 (2010).

45. Son, H.-J. & Song, K. Bin. Antimicrobial Activity of Flaxseed Meal Extract against Escherichia coli O157: H7 and Staphylococcus aureus Inoculated on Red Mustard. J. Microbiol. Biotechnol. 27, 67--71 (2017).

46. Tian, J. et al. Antifungal cyclic peptides from Psammosilene tunicoides. J. Nat. Prod. 73, 1987–1992 (2010).

47. Pinto, M. E. F. et al. Ribifolin, an orbitide from jatropha ribifolia, and its potential antimalarial activity. J. Nat. Prod. 78, 374–380 (2015).

48. Röttig, M. et al. NRPSpredictor2 - A web server for predicting NRPS adenylation domain specificity. Nucleic Acids Res. 39, W362–367 (2011).

49. Mingxun, W. et al. Assembling the Community-Scale Discoverable Human Proteome. Cell Syst. Press

50. Keller, B. O., Sui, J., Young, A. B. & Whittal, R. M. Interferences and contaminants encountered in modern mass spectrometry. Analytica Chimica Acta 627, 71–81 (2008).

51. Arnison, P. G. et al. Ribosomally synthesized and post-translationally modified peptide natural products: overview and recommendations for a universal nomenclature. Nat. Prod. Rep. 30, 108–160 (2013).

## SUPPLEMENTARY REFERENCES

1. Mohimani, H. et al. Dereplication of peptidic natural products through database search of mass spectra. Nat. Chem. Biol. 13, 30–37 (2017).

2. Röttig, M. et al. NRPSpredictor2 - A web server for predicting NRPS adenylation domain specificity. Nucleic Acids Res. 39, W362–367 (2011).

3. Frank, A. M. et al. Spectral archives: extending spectral libraries to analyze both identified and unidentified spectra. Nat. Methods 8, 587–591 (2011).

4. Mingxun, W. et al. Assembling the Community-Scale Discoverable Human Proteome. Cell Syst. Press

5. Keller, B. O., Sui, J., Young, A. B. & Whittal, R. M. Interferences and contaminants encountered in modern mass spectrometry. Analytica Chimica Acta 627, 71–81 (2008).

6. Arnison, P. G. et al. Ribosomally synthesized and post-translationally modified peptide natural products: overview and recommendations for a universal nomenclature. Nat. Prod. Rep. 30, 108–160 (2013).

7. Kavan, D., Kuzma, M., Lemr, K., Schug, K. A. & Havlicek, V. CYCLONE - A utility for de novo sequencing of microbial cyclic peptides. Journal of the American Society for Mass Spectrometry 24, 1177–1184 (2013).

8. Gurevich, A. et al. Increased diversity of peptidic natural products revealed by modification-tolerant database search of mass spectra. Nat. Microbiol. 3, 319–327 (2018).

